# Deep sequencing of natural and experimental populations of *Drosophila melanogaster* reveals biases in the spectrum of new mutations

**DOI:** 10.1101/095182

**Authors:** Zoe June Assaf, Susanne Tilk, Jane Park, Mark L. Siegal, Dmitri A. Petrov

## Abstract

Mutations provide the raw material of evolution, and thus our ability to study evolution depends fundamentally on whether we have precise measurements of mutational rates and patterns. Here we explore the rates and patterns of mutations using i) *de novo* mutations from *Drosophila melanogaster* mutation accumulation lines and ii) polymorphisms segregating at extremely low frequencies. The first, mutation accumulation (MA) lines, are the product of maintaining flies in tiny populations for many generations, therefore rendering natural selection ineffective and allowing new mutations to accrue in the genome. In addition to generating a novel dataset of sequenced MA lines, we perform a meta-analysis of all published MA studies in *D. melanogaster*, which allows more precise estimates of mutational patterns across the genome. In the second half of this work, we identify polymorphisms segregating at extremely low frequencies using several publicly available population genomic data sets from natural populations of *D. melanogaster*. Extremely rare polymorphisms are difficult to detect with high confidence due to the problem of distinguishing them from sequencing error, however a dataset of true rare polymorphisms would allow the quantification of mutational patterns. This is due to the fact that rare polymorphisms, much like *de novo* mutations, are on average younger and also relatively unaffected by the filter of natural selection. We identify a high quality set of ~70,000 rare polymorphisms, fully validated with resequencing, and use this dataset to measure mutational patterns in the genome. This includes identifying a high rate of multi-nucleotide mutation events at both short (~5bp) and long (~1kb) genomic distances, showing that mutation drives GC content lower in already GC-poor regions, and finding that the context-dependency of the mutation spectrum predicts long-term evolutionary patterns at four-fold synonymous sites. We also show that *de novo* mutations from independent mutation accumulation experiments display similar patterns of single nucleotide mutation, and match well the patterns of mutation found in natural populations.

## Introduction

Mutation is the ultimate driver of genetic diversity. All genetic differences found both within and between species originated in the mutational process, and then survived the stochastic and selective forces acting on every allele’s frequency dynamics. As a result, the patterns in population genetic and evolutionary data are a combined result of both selective and mutational forces. Studies of natural selection thus depend fundamentally on whether we can correct for the confounding factors of mutational biases.

Our ability to study mutation is severely limited both because mutation rates are generally extremely low, and because a substantial fraction of new mutations are deleterious and thus purged from populations by purifying selection. These two problems can, at first glance, be overcome with divergence-based measurements, in which rates of substitution within nonfunctional genomic regions are calculated across taxa [1–3]. Here the use of vast phylogenetic timescales allows one to observe large numbers of even very rare mutational events, while the use of neutral sequences eliminates the confounding effects of natural selection. Unfortunately, these divergence-based methods are compromised by the necessity of identifying large numbers of regions that behave truly neutrally [4–6], and by the fact that even mutations residing in truly neutral regions can be subject to the selection-like force of biased gene conversion [7,8] or to selection on genome properties such as GC content or genome size [9–11]. Furthermore, divergence based methods produce long term averages for the mutational spectrum, and thus may be sensitive to variation in life history traits such as generation time [12] and may obscure mutation rate variation through time [13].

The ideal approach for the study of mutational processes is to identify truly new mutations. The optimal study would capture *de novo* mutations via sequencing of sets of parents and offspring, and implement this strategy in an enormous sample in order to overcome both inter-individual mutation rate variation and the small number of events that occur per individual. The advent of next-generation sequencing has made this possible, however it is still quite expensive and only recently beginning to be realized, primarily in humans [14]. Thus, in order to measure mutational rates and biases in organisms that do not receive the same level of funding support as humans, or in order to survey human mutational rates and patterns across diverse populations on a reasonable budget and timescale, we must use alternative approaches.

Two such alternative approaches include mutation accumulation (MA) experiments in model organisms [15], and a more recently proposed method in which very rare polymorphisms in natural populations are used as a proxy for new mutations [16,17]. These approaches have complementary strengths and weaknesses.

The MA approach is implemented by maintaining an organism in a population so small (*N* ~ 1 – 2) that selection is ineffective (*Ns* ≪ 1) for even strongly deleterious mutations, thus allowing (nonlethal) mutations to be passed along through the generations [15]. This experiment results in an accumulation of new mutations on the chromosomes, which, with enough individuals or enough generations, turns the relatively infrequent event of mutation into an observable process. A testament to the utility of this method is perhaps the sheer number of organisms in which MA has been deployed: *Arabidopsis* (*thaliana, floriosa, and douglasiana), Caenorhabditis (elegans and briggsae), Chlamydomonas reinhardtii, E. coli, O. myriophila, S. cerevisiae, VSV, ϕ6 virus* and *Drosophila melanogaster* [15]. MA experiments have the benefit of allowing researchers to precisely measure both mutational rates and mutational patterns in a controlled laboratory environment. However therein also lie the weaknesses of MA approaches: i) the controlled environment and the use of lab strains makes it possible that mutational rates and patterns are not representative of what occurs in natural populations, ii) the accumulation of deleterious mutations during the course of the MA experiments may itself change the spectrum and rate of new mutations, and iii) the MA experiments can be extremely laborious and thus limited in the sheer number of events they are likely to generate.

The second approach to studying the mutational spectrum uses very rare polymorphisms as a proxy for new mutations. The rationale of this approach is that very rare polymorphisms are younger on average and have frequency dynamics dominated by stochastic forces rather than by natural selection or biased gene conversion [16,18]. An ideal example of such a mutation would be a singleton, or in other words, a mutation present on just a single chromosome in a population (or population sample). Such a mutation was likely generated in the germline of the previous generation and thus private to the current individual within which it resides, and it’s frequency relatively unaffected by natural selection. Thus, the expectation is that as we look at polymorphisms at lower and lower population frequencies, we should observe classes of genetic variants with probabilities that become primarily determined by mutational biases. The principal advantages of this approach are that we can study mutational patterns as they occur in nature, and that we can obtain very large numbers of events economically. This strategy also has its own drawbacks, however, including: i) the inability to directly measure mutation rates, ii) the challenge of ensuring that selection has had minimal effect on the patterns of mutations, and iii) the difficulty of distinguishing true variants from sequencing and alignment errors. The last problem is particularly daunting. As Achaz (2008) noted, increasing the number of individuals in the sample does not help - the number of errors and the number of true variants both scale linearly with sequence length, but an increase in sample size causes the number of true variants to scale only logarithmically while the number of errors still scales linearly [19]. Thus, in the pursuit of a deep catalogue of genetic variation in natural populations, it is quite possible that as more and more individuals are sequenced, we will be adding disproportionately more errors than real polymorphisms.

In this study we leverage the opportunity to combine the MA strategy with the rare polymorphism approach, thus avoiding the drawbacks of either method in isolation while also benefiting from the strengths of both. Mutations generated in the controlled environment of the laboratory provide an exciting opportunity to directly measure mutation rates, while also generating a neutral expectation which can be compared with a much larger dataset of rare polymorphisms segregating in natural populations. This combined approach allows a nuanced characterization of the mutational spectra, whether generated in the laboratory or in natural populations, and creates deep enough data that mutational patterns can be studied across the genome in fine detail.

The fruit fly *Drosophila melanogaster*, as both a model organism and a species with a large number of sequenced natural isolates, provides an excellent opportunity to implement this integration of approaches. In the first half of this study, we combine results from five MA experiments, including our own novel dataset, to arrive at a set of 2,141 *de novo* mutations which were generated in the laboratory. Then, in the second half of this study, we use 3 large datasets of sequenced natural populations in order to extract a large set (~ 70, 000) of high quality rare (< 0.1%) polymorphisms which, unlike other studies, have been fully validated via resequencing. We use these data to validate both MA and rare polymorphism approaches for the study of new mutations, to provide the most precise estimate to date of the rate and patterns of point mutations, and to detect substantial neighbor-dependency of mutation in *D. melanogaster*.

## Results

### *De novo* mutations identified in MA experiments

While mutation accumulation studies within the fruit fly community have a history dating back to the early 20th century [20], it’s only in the last several years that researchers have been able to directly measure the single base pair mutation rate using sequencing methods [21–24]. A number of different experimental designs have been used, and perhaps the most marked difference between them is the choice of whether to use the homozygous vs heterozygous MA strategy. Homozygous MA is essentially inbreeding in a small population (*N* ~ 2) which therefore forces new mutations to eventually be homozygosed (Figure 1A, right). The heterozygous MA experiment uses a cross scheme which instead passes the MA chromosome in a heterozygous state through a single male in every generation (Figure 1A, left). These different approaches result in different amounts of selection that occurs against recessive strongly deleterious mutations, a class of mutational events that are both critical to fitness and common in many natural populations [25]. At the outset of this study there were two prior publications that sequenced MA lines, both of which used a homozygous MA strategy [21,22], and thus we sought to contribute a novel dataset generated via heterozygous mutation accumulation. When MA experiments were first invented the heterozygous strategy was often implemented with the inversion-rich balancer chromosomes, that may be prone to distortions in homology-directed repair processes. We thus implemented a more conservative approach of using un-inverted chromosomes carrying recessive markers (see Methods), which more closely mimic nature. Since we began our experiments, there have been two more recent datasets published [23,24]. In this work we present our own novel dataset from a heterozygous MA experiment, and then generate the first meta-dataset of all MA studies in the fruit fly yet published, with which we precisely estimate mutational patterns both across experiments and within the combined dataset.

**Figure 1:**
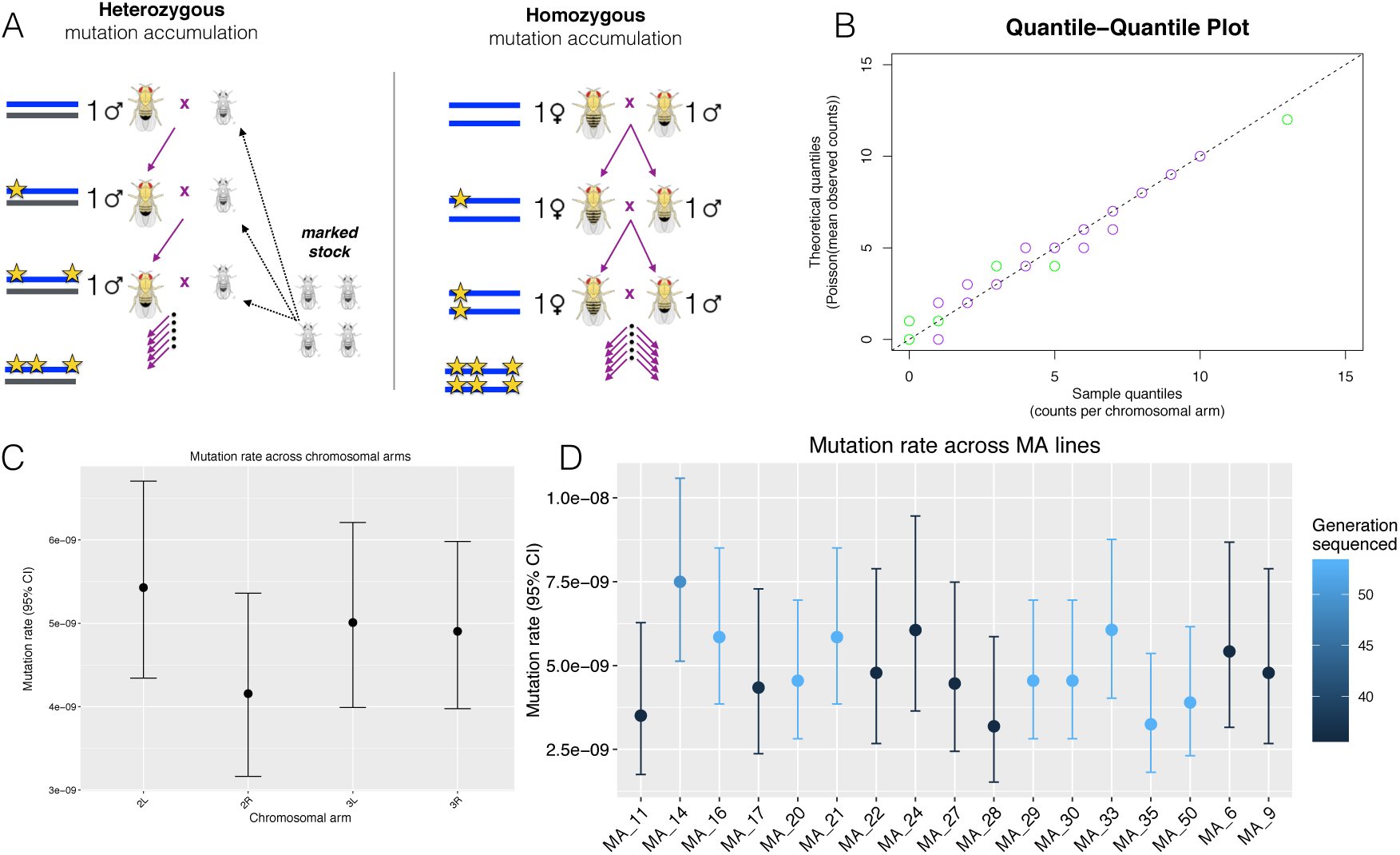
A summary of the experimental design and results for the single base pair mutation rate in this study. (A) Diagram depicting the general cross schemes used in heterozygous (left) and homozygous (right) mutation accumulation, where this study used the heterozygous design. (B) QQ plot of the quantiles of the mutation counts on each chromosome arm of each strain, plotted against the quantiles of a Poisson distribution with mean taken from the mean counts in the MA experiment, where color indicates the generation sequenced (green = generation 36, purple = generation 53). (C) Mutation rates estimated for each chromosomal arm (Pearson’s Chi-squared test of independence, X-squared = 2.55, df = 3, p-value = 0.47). (D) Mutation rates estimated for each strain, where color indicates the generation sequenced (Pearson’s Chi-squared test of independence, X-squared = 7.99, df = 14, p-value = 0.89).

#### A new dataset of 325 point mutations

Briefly, our new heterozygous MA dataset was generating using the following experimental and sequencing pipelines: 17 independent lines of *D. melanogaster* were allowed to accumulate mutations for 36-53 generations in a heterozygous state (Figure 1A, left). Each line was then sequenced to ~20-25X (sample coverage map can be seen in Figure S1), sequencing reads processed (trimmed, mapped to release 5.57 of the flybase reference, filtered for duplicates, and realigned around indels), and variants called with a combination of GATK and Varscan. A variant was considered a de novo mutation if it was called with high confidence in one strain and simultaneously never present on more than a single sequencing read in either the ancestral strains or any other MA line. In total 325 new mutations were identified, of which 30 were randomly chose for visual confirmation in a pileup file and an additional 30 were randomly chosen for PCR/Sanger sequencing. We successfully validated 29 of the 30 that were Sanger sequenced, giving a ~ 3% error rate, although we note that upon visual inspection of the single unconfirmed mutation, we verified that both the original genotype call and the resequence data to be of very high quality, and consequently we suspect the PCR primers may have inadvertently been haplotype-specific and thus amplified the non-MA chromosome. See Methods for additional details of pipeline, and Tables S1, S2, and S3 for details of the 17 MA strains, 325 *de novo* mutations identified, and 30 mutations Sanger sequenced.

The 325 *de novo* mutations identified in this study reveal a notably consistent mutation rate across strains, chromosomes, and time. Plotting the quantiles of mutation counts on chromosomal arms against the quantiles of a Poisson distribution (with a mean equal to the sample mean) gives a markedly linear relationship (Figure 1B), affirming that mutation counts are indeed Poisson distributed. Consistent with previous findings [21,22], we find no significant difference in mutation counts across the major chromosomal arms (Figure 1C, *χ*^2^ test p-value = 0.47). Additionally, we find no significant difference in mutation rates between generations 36 and 53 (Poisson exact p=0.76, Figure S2), although it is likely we were underpowered to test for small variations in rates across generations. Given the presence of mutator lines in previous MA experiments [22,23] it was also important to test for variation in the total mutation rate across strains, and we find no evidence for any variation in mutation rates across strains (Figure 1D, *χ*^2^ test p-value = 0.89). Finally, in our experiment we found a single base pair mutation rate of 4.9e-9 per generation (95% CI 4.4 – 5.5e-9).

#### Five MA experiments, different single base pair mutation rates

We next compared and combined our dataset with data from four previously published experiments [21–24]. Across all five experiments there are in total 163 lines which went through between 36 and 262 generations, however of those there were five lines (four from [22], one from [23]) with elevated mutation rates, and thus we reduced the set to the 158 non-mutator strains for our combined analyses. We restrict our comparisons to the major autosomes only (chromosomes 2 and 3), as these were the chromosomes used across all five MA experiments and upon which most mutations resided (note that the inbreeding MA strategy does allow accumulation on chromosomes X and 4 as well). Additionally, we masked repeats in all data sets (see Methods for additional details of obtaining and processing these data). This procedure of filtering out mutator lines, chromosomes X and 4, and repeat regions, reduces the total number of mutations across all experiments from 3,187 to 2,141. We work with the dataset of 2,141 mutations when doing comparisons across experiments, however we also make the entire dataset available for download.

A comparison of experiments and single base pair mutation rates can be found in Table 1. Our mutation rate is significantly higher than that reported by the homozygous MA accumulation studies of both Keightley et. al. (Poisson exact p=2e-4) and Schrider et. al. (Poisson exact p=3e-6), significantly lower than that reported by the heterozygous MA of Sharp et. al. (Poisson exact p=1.5e-3), and not significantly different from that reported by Huang et. al. (Poisson exact p=0.35) (see Methods and Table S4 for additional details). Overall, the experimental designs which used heterozygous accumulations, fewer generations, and newer sequencing technologies, tended to have higher mutation rate estimates, emphasizing that experimental design is an important consideration in MA studies.

**Table 1:** A summary of the five mutation accumulation experiments analyzed here, which in addition to ours include references [21–24]. The combined data we work with in this study consists of the ‘filtered’ set of mutations, a subset of the original data available from refs. [21–24], in which we use major autosomes, non-repetitive regions, and nonmutator lines only (mutators include line 19 from Huang et. al., and lines from ancestor 33 in Schrider et. al.). The mutation rates and 95% confidence intervals are the rates and intervals provided by the published studies. Note that Huang et. al. reported the median mutation rate only, thus we approximated the mutator line’s rate from a figure in the paper (Fig7) and we re-constructed a reasonable CI given the data. Our mutation rate (4.9e-09) is significantly higher than the rates reported by both Keightley et. al. (Poisson exact p=2.03e-04) and Schrider et. al. (Poisson exact p=3.4e-06), significantly lower than that reported by Sharp et. al. (Poisson exact p=1.5e-03), and not significantly different from that reported by Huang et. al. (Poisson exact p=0.35).

| Study | MA method | # Lines | Generations per line (approx.) | Mutation count (chr 2/3) | Mutation rate |
| --- | --- | --- | --- | --- | --- |
| **MA combined** | - | 158 | - | 2259 | - |
| **Assaf 2016** | heterozygous | 17 | 45 | 325 | 4.90E-09 (4.4-5.5e-9) |
| **Sharp 2016** | heterozygous | 112 | 60 | 786 | 6.03E-09 (5.6-6.5e-9) |
| **Huang 2016** | hybrid | 22 | 52 | 823 | 5.21E-09 (unavailable) |
| **Schrider 2013** | homozygous | 4 | 145 | 175 | 3.27E-09 (2.9-3.7e-09) |
| **Keightley 2009** | homozygous | 3 | 262 | 150 | 3.07E-09 (2.6-3.6e-09) |

#### The neutral expectation is reached in all five experiments

We can next look at functional regions in the genome in order to test whether the mutational spectra across the five MA experiments are truly unbiased by natural selection. While polymorphisms within functionally important sites (i.e. nonsynonymous or nonsense, which tend to be deleterious) typically do not reach high frequencies in natural populations, we do expect that *de novo* mutations occurring in coding regions should cause a nonsynonymous change ~75% of the time, and cause a nonsense change ~4% of the time. As can be seen in Figure 2A-B, the five mutation accumulation experiments do indeed exhibit the expected fractions of 75% nonsynonymous and 4% nonsense (no significant difference between experiments, *χ*^2^ test p=0.03 and p=0.69 for nonsynonymous and nonsense respectively), although note that the counts are not high enough to give a well-defined point estimate for the nonsense fraction.

**Figure 2:**
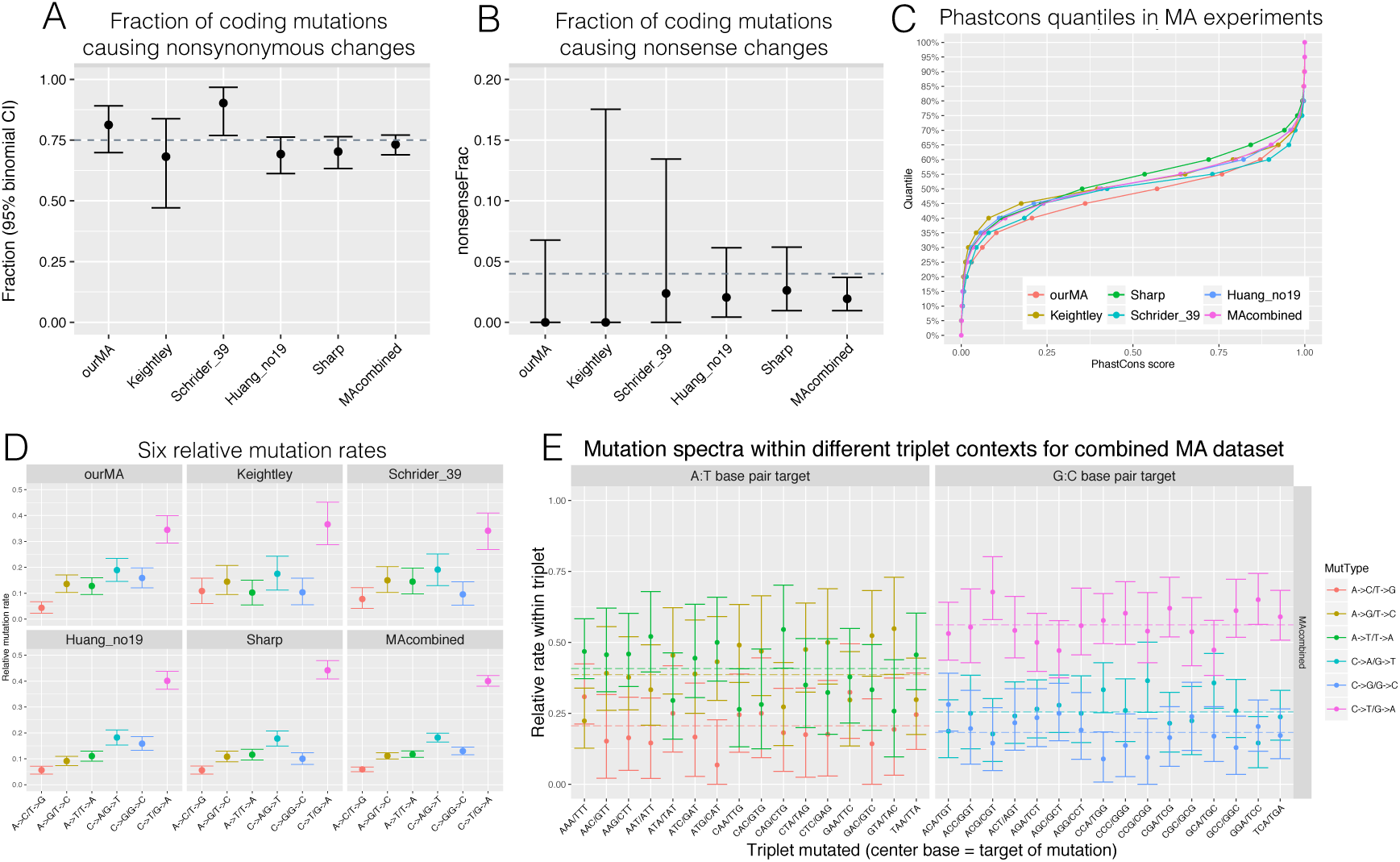
A summary of comparisons conducted between the five different MA experiments, including (A) the six relative mutation rates (i.e. sum to 1) (B) the fraction of coding mutations which cause nonsynonymous changes, where the dotted line indicates the neutral expectation of 75%, (C) the fraction of coding mutations which cause nonsense changes, where the dotted line indicates the neutral expectation of 4%, and (D) the empirical cumulative distribution for phastCons scores within each MA experiment.

Another method to test if the MA mutations were generated in the absence of selection is to classify all sites in the genome by their conservation status across phylogenetic timescales, and then ask whether the distribution of mutations from the MA experiments meet the neutral expectation. To do this we employed the publicly available phastCons scores for the *D. melanogaster* reference genome, which are a measure of evolutionary conservation across twelve Drosophila species, mosquito, honeybee, and the red flour beetle [26]. Indeed, as we can see in Figure 2C, the distribution of phastCons scores are similar across MA experiments, and not significantly different from the neutral expectation as given by the distribution of scores in the reference genome (bootstrap KS test p=0.31).

#### Mutational spectra are comparable across all five experiments

Mutations can be collapsed into six basic types, and, after scaling for the *D. melanogaster* genome GC content of 43%, their relative mutation rates calculated (Figure 2D, Table S5). We find that these six relative rates are indeed significantly different across experiments (*χ*^2^ test, p=0.003), however the p-value is not very low. Indeed, comparisons of various subsets of the data gave generally insignificant differences, such that no specific mutation type nor specific experiment was found to be driving the variation (tests of single mutation types or single experiments against the sum of the rest gave *χ*^2^ test corrected p-values>0.01). We next combined the datasets to generate a more precise estimate of the six relative rates (bottom right Figure 2D, Table S5). The *C* → *T*/*G* → *A* mutation type, in addition to occurring at comparable relative rates across experiments (*χ*^2^ test p=0.15), is by far the most common mutation (0.4 relative rate, 95% CI 0.38-0.42). In *D. melanogaster* it occurs at ~7X the rate of the least common *A* → *C*/*T* → *G* mutation type. This is despite the paucity of cytosine methylation in *D. melanogaster* [27], which in humans, for example, drives a relative rate of *C* → *T*/*G* → *A* of ~0.48 (which is ~ 11*X* the least common mutation type in humans) [14]. Our finding is consistent with previous work in *Drosophila* and recent work in yeast [28–30] showing an elevated mutation rate at cytosines despite minimal or no methylation in the genome, suggesting that the sensitivity of cytosines to mutation may be a general feature of cytosines in a cellular context.

Using the counts and relative rates of the six mutation types we can now look at transition:transversion ratios and GC equilibrium, which we find to be similar across experiments. When considering the number of transition mutations (2 possible types) and transversion mutations (4 possible types), we find transition:transversion ratios that do not vary significantly across experiments (G test of independence, p=0.21), and which for the combined dataset reaches a ratio of 2:1 (95% CI 1.9-2.2) (Table 2). Next, by considering mutations which change the GC content of the mutated base pair, we can ask if the mutational process tends to drive the genome towards A:T base pairs or G:C base pairs by calculating the GC equilibrium. The GC equilibrium has only been reported by one MA experiment [21], however we can systematically quantify it across all five experiments (Table 2, significantly different between experiments, G test p=0.01). Interestingly, while the oldest MA paper reported a GC equilibrium of 30% (with a large CI of 24-40% [21]), we find that newer studies have consistently found a lower value. For the combined dataset the GC equilibrium reaches ~23% (95% CI 0.21-0.25). In contrast, the *D. melanogaster* genome has an actual GC content of 43%, and thus the lower GC equilibrium found here emphasizes the importance of non-neutral processes in driving the genome GC content higher [8].

**Table 2:**
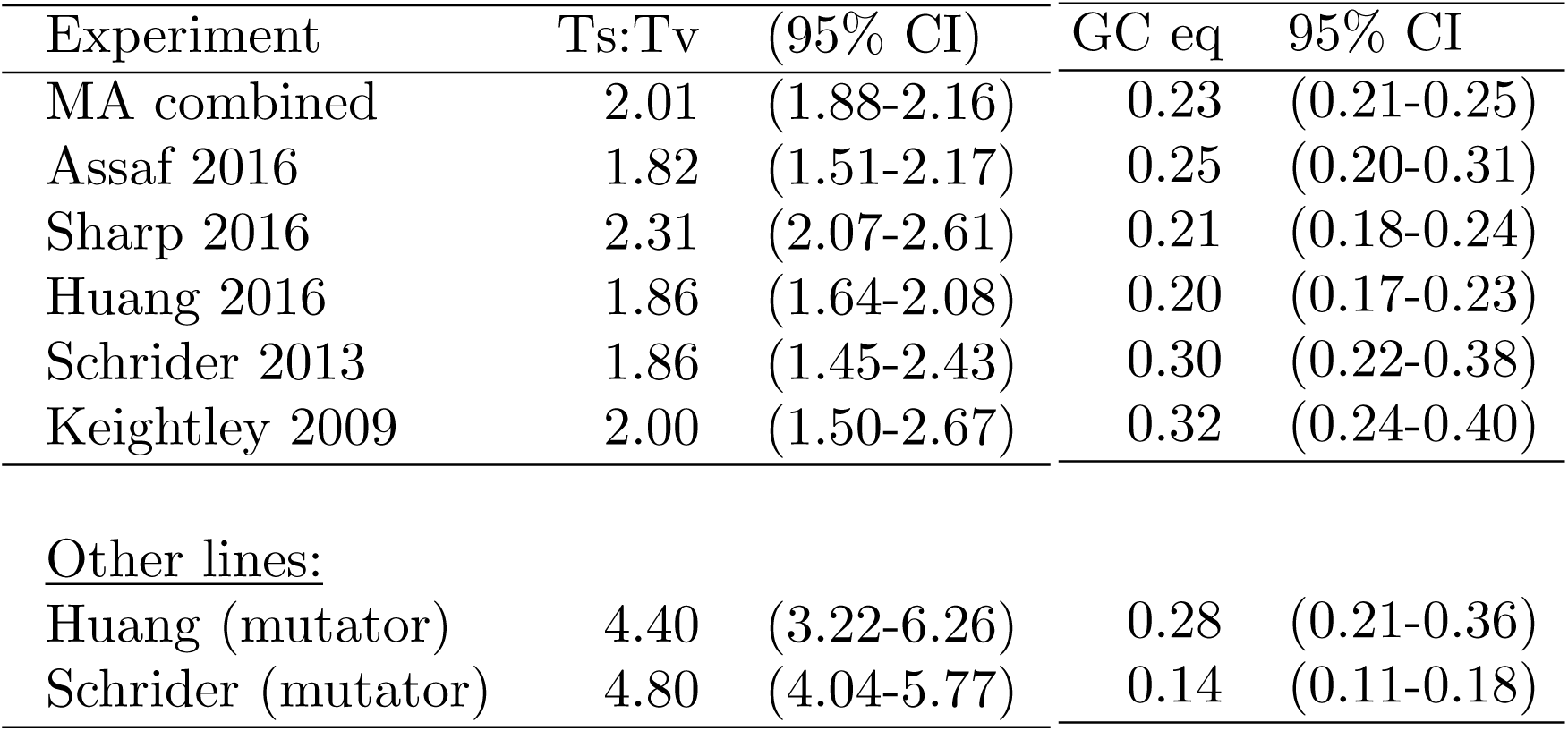
A summary of the transition:transversion ratios across experiments, which are not significantly different (G test of independence, p=0.21), and which for the combined set is ~2:1. The 95% confidence intervals were calculated via 1000 bootstraps of raw counts. Ts:Tv ratio is calculated as: (count of transition mutations) / ((count of transversion mutations)/2).

Lastly, we can also test for neighbor-dependent variation in the mutation spectrum. There is evidence in some organisms that single base pair mutation rates can vary depending on the neighboring base pair context [17, 24, 29]. To test this in *D. melanogaster* using our combined MA dataset we considered triplet contexts, in which the center base is mutated. All possible triplets were collapsed into their forward/reverse sequence, and then we quantified within each triplet the relative rates of the three mutation types that can occur at the center base pair of the triplet (e.g. CAG/CTG triplet can have A→T/T→A, A→C/T→G, or A→G/T→C mutations). In contrast to quantifying the total mutation rate within each triplet, measuring the relative rates within each triplet provides an internal control for the triplet content in the genome, which will vary across MA publications depending on which masks were applied to the reference genome (information that is not consistently documented across publications). Using the combined set of 2,141 *de novo* mutations from the five MA experiments, we do in fact find heterogeneity in the mutation spectrum across triplet contexts (Figure 2E, G test of independence, p=0.008 and p=0.007 for GC and AT base pairs respectively). However, 2,141 mutational events is not a large enough dataset to detect whether any particular triplet is driving the heterogeneity (G test, corrected p values > 0.01, Table S6). Thus, despite compiling the largest *Drosophila* dataset of *de novo* mutations yet available, it would be desirable to generate an even larger number of mutational events.

### Rare polymorphisms identified in natural populations

#### Identification and validation of rare polymorphisms

As a proxy for new mutations, we seek to identify a class of ultra-low-frequency polymorphisms. To this purpose, we use three publicly available datasets (Table 3) and employ the method depicted in Figure 3A and briefly described here: i) We first downloaded 621 genomes from the Drosophila Genome Nexus (DGN) [31], which represent predominantly monoallelic genomes from 35 populations across 3 continents that underwent the same iterative mapping pipeline before variant calling. These data represent an extremely high quality set of genotype calls, and thus we identified all genetic variants with which we will be working using these data (Step 1 in Figure 3A, Methods). ii) We next leveraged the availability of pooled sequencing data generated by our and collaborating labs, which collectively represent >17,000X coverage of >4,000 genomes from across the east coast of the USA and Europe [32] [plus additional unpublished data]. Pooled sequencing data is not ideal for rare variant identification, due to the difficulty of distinguishing a rare polymorphism from a sequencing error [33]. In order to circumvent this, we used the set of high quality DGN singletons identified in Step 1, i.e. those at 1/621 frequency, and filtered them down by removing those that appeared in the pooled sequence data. This allowed the identification of DGN singletons at a frequency that is an order of magnitude lower (frequency ~1/5000 = 0.0002) (Step 2 in Figure 3A, Methods). iii) Finally, the third public resource we leveraged is resequence data made available by the DGRP and DPGP1 projects, which independently resequenced 29 strains (present in the DGN) using 454 and illumina sequencing [unpublished but available online]. Given that any pipeline which filters down to polymorphisms unique to a single genome is likely to be enriching for sequencing error (i.e. a polymorphism segregating in multiple individuals or multiple datasets is less likely to be an error), we further validated our rare DGN variants by requiring an identical genotype call to be made in the resequence data (Step 3 in Figure 3A, Methods). This procedure does reduce the number of polymorphisms down to only those which appeared in the 29 resequenced strains (Table 4), however this dataset is of extremely high quality.

**Figure 3:**
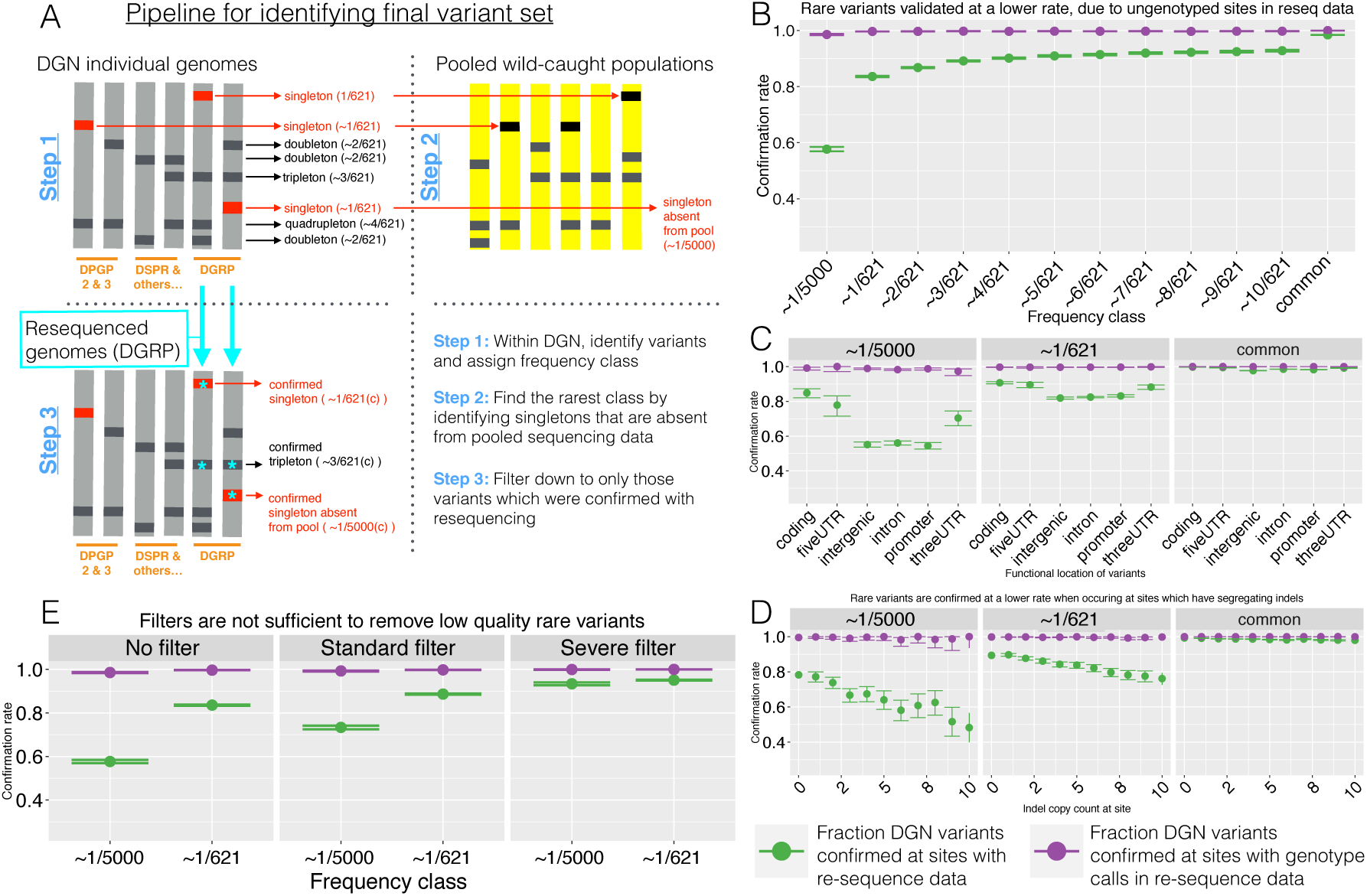
Description of the data pipeline and quality used in the identification of low frequency polymorphisms. (A) Diagram of Steps 1 thru 3 in identifying a high quality set of rare polymorphisms, (B)-(E) depict confirmation rate where **green** indicates the fraction of genotype calls within the DGN data (identified in Step 1) which were confirmed in the resequence data (Step 3), and **purple** indicates the fraction of genotype calls within the DGN data which were not disconfirmed (i.e. using polymorphisms for which a genotype call exists in both the DGN and resequence data to measure the confirmation rate). The confirmation rate is depicted as a function of (B) frequency, (C) of genomic location, (D) of segregating indel copy number, and (E) of the filters applied to the dataset. For (E) the filters include no filters, standard filters (QD > 2, QUAL > 20, 3>DP>100), and severe filters (QD > 3, QUAL > 55, 9>DP>100, and requiring that each genotype call is at a site for which 85% of individuals have a genotype call and for which the total copy number of indels segregating in other individuals is ≤10).

**Table 3:** A summary of the deep sequence data from natural populations of *D. melanogaster* used in this study for the identification of low frequency polymorphisms.

| Data source | Citation | Description |
| --- | --- | --- |
| DGN genomes | Lack et. al. 2015 | High coverage individual genomes. All genomes went through the same iterative mapping pipeline, where raw sequence reads were sourced from the study's authors and several additional resources including DGRP, DPGP, DPGP3, and DSPR. The sequenced flies were derived from isofemale inbred lines which were sequenced either is or after generating haploid embryos. |
| Pooled wild populations | Bergland et. al. 2014 | High coverage sequencing of pooled wild caught flies. Data acquired via the NESCENT project (unpublished), which consists of >17000X coverage of >4000 genomes from ~30 populations sequenced in pools. |
| Resequencing data | unpublished | High coverage resequencing of individual genomes (which are included in the DGN). Consists of 29 of the inbred strains from the DGRP project, sequenced independently by the DPGP1 project (separate DNA extraction and library prep, sequenced on the illumina platform) and/or sequenced by the DGRP project (same DNA extraction, separate library prep, sequenced on the 454 platform). |

**Table 4:**
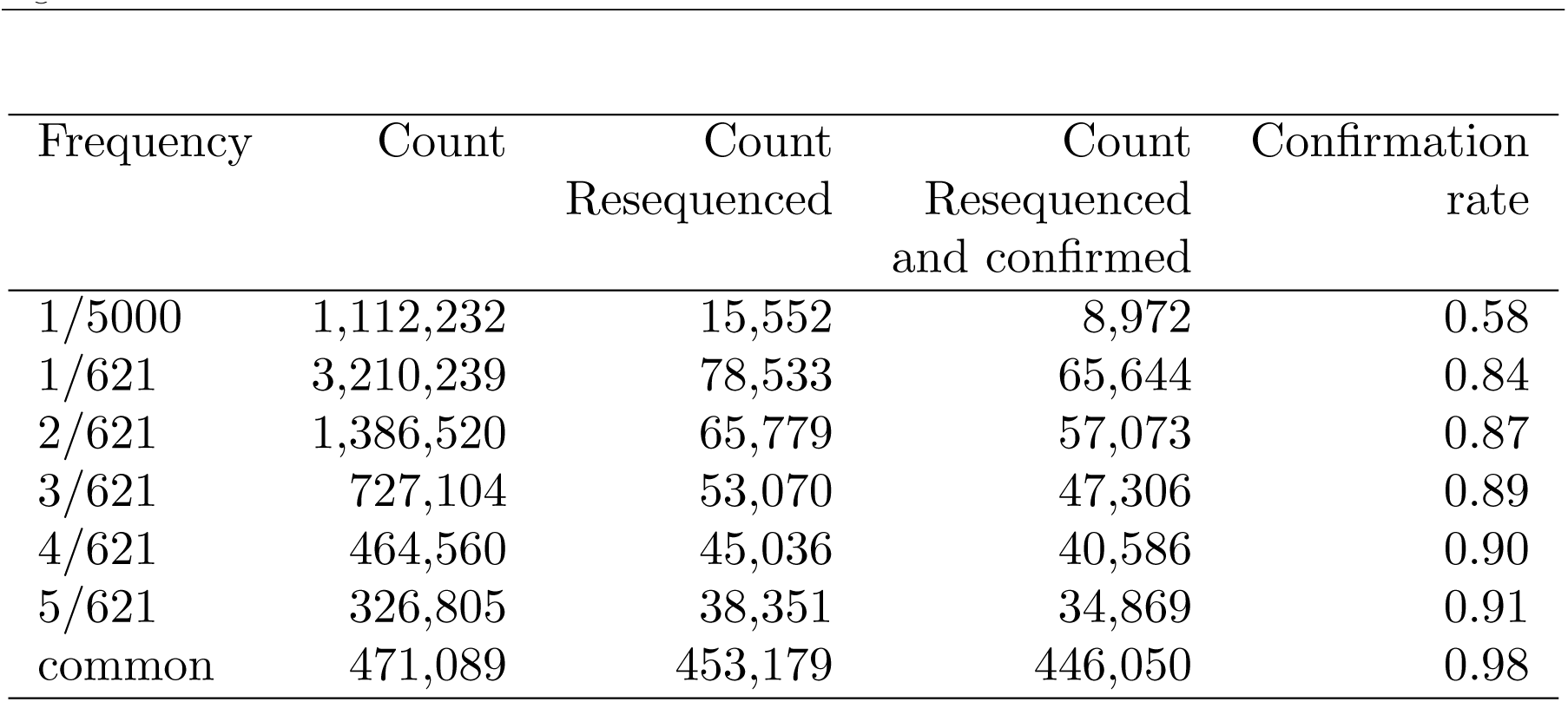
The count of polymorphisms for each frequency class when looking across the entire dataset (second column), the resequenced dataset (third column), and the dataset of variants resequenced and confirmed (fourth column). It can be seen that the confirmation rate decreases with decreasing polymorphism frequency.

| Frequency | Count | Count<br>Resequenced | Count<br>Resequenced<br>and confirmed | Confirmation<br>rate |
| --- | --- | --- | --- | --- |
| 1/5000 | 1,112,232 | 15,552 | 8,972 | 0.58 |
| 1/621 | 3,210,239 | 78,533 | 65,644 | 0.84 |
| 2/621 | 1,386,520 | 65,779 | 57,073 | 0.87 |
| 3/621 | 727,104 | 53,070 | 47,306 | 0.89 |
| 4/621 | 464,560 | 45,036 | 40,586 | 0.90 |
| 5/621 | 326,805 | 38,351 | 34,869 | 0.91 |
| common | 471,089 | 453,179 | 446,050 | 0.98 |

This procedure confirms that, indeed, the proportion of artifactual variants increases as their frequency decreases (Figure 3B-D), and as a consequence it is absolutely critical to validate rare variants using resequence data. We find that while common DGN variants are confirmed in the resequence data at a rate close to ~100%, the rarest DGN variants, at frequency ~1/5000, have a much lower confirmation rate in the resequence data of <60% (Figure 3B, green points, Table 4). We find this to be primarily driven by sites that are not genotyped at all during resequencing, due to the fact that if we look at only DGN variants which were successfully genotyped in the resequence data, we find the confirmation rate rises back to ~100% (Figure 3B, purple points). This low confirmation rate for rare polymorphisms appears to be largely driven by low complexity and indel-rich regions (Figure 3C-D). As can be seen in Figure 3C, rarer DGN polymorphisms have a lower confirmation rate in intronic and intergenic regions (Figure 3C, left and center panels), an effect which is negligible for common variants (Figure 3C, right panel), and which disappears when looking at only DGN variants with a genotype call in the resequence data (Figure 3C, purple points). As can be seen in Figure 3D, despite masking each DGN genome for indels before calling variants in that individual (Methods), rare DGN variants have a low confirmation rate within sites at which indels are segregating in other individuals (Figure 3D, left and center), an effect that is exacerbated as the indel frequency (i.e. copy count) increases in the DGN population. This effect is again negligible for common variants (Figure 3D, right panel), and disappears when looking at only DGN variants with a genotype call in the resequence data (Figure 3D, purple points).

These results beg the question of whether more severe filtering can approximate the quality-control achieved with resequencing, i.e. does the confirmation rate recover if we filter down to DGN genotype calls which have better scores for metrics like depth and quality score? To ask this we measured confirmation rate after implementing standard filters used by most researchers (site has QUAL > 20, QD > 2, 3>DP>100 (note mean depth of data is ~ 25X)), as well as much more severe filters (site has QD > 3, QUAL > 55, 9>DP>100, genotype data in ≥ 85% of samples, indels in ≤ 10 samples). As can be seen in Figure 3E, the standard filters used by many researchers give rare polymorphisms that still have a confirmation rate as low as ~70%, and while severe filters do better at a ~90% confirmation rate, this rate is not ideal given that a tenth of the data may still be error-prone. These results make sense if we look at the distribution of DP and QUAL scores within the DGN data for sites that are confirmed, disconfirmed, and ungenotyped in the resequence data. While the distributions of scores are significantly different, they also overlap (Figure S3), and thus even severe filters are likely to let through variant calls that may be errors. These results emphasize the importance of resequencing genomes when working with polymorphisms that are segregating at low frequency. The final count of rare polymorphisms that we will use in this study, all confirmed via resequencing, can be seen in Table 4.

The reduced confirmation rate within introns and intergenic regions, as well as near indels, emphasizes the difficulty of validating rare polymorphisms within low-complexity regions. It has been observed before that artifactual variant calls in high-coverage sequencing samples are largely driven by alignment errors [34]. This is potentially worrisome when testing for mutational biases in the genome, because some tests rely inherently on our ability to ‘count the reference’, meaning accurately quantifying in the reference genome the relative proportions of sequence contexts in which we might be interested. For example, it is known that regions with higher GC content tend to have higher coverage [35], and thus potentially a higher discovery rate of genetic variants. Thus, even when we are confident our genetic variants are real, when working with rare polymorphisms we must be careful not to confound intrinsic rates of detection with intrinsic rates of mutation. For this reason, when using metrics which rely on quantifying relative proportions of sequence context in the reference genome (e.g. GC equilibrium, six relative rates, neighbor-dependency), we prefer to work with higher complexity zones such as coding regions (as in Figure 4), or alternatively re-frame the metric in such a way as to not be sensitive to this factor (as in Figure 2E). We note that a similar approach is used in other studies of context-dependent mutational patterns [17], and that while our results are robust to this choice we consider it to be the more conservative approach.

**Figure 4:**
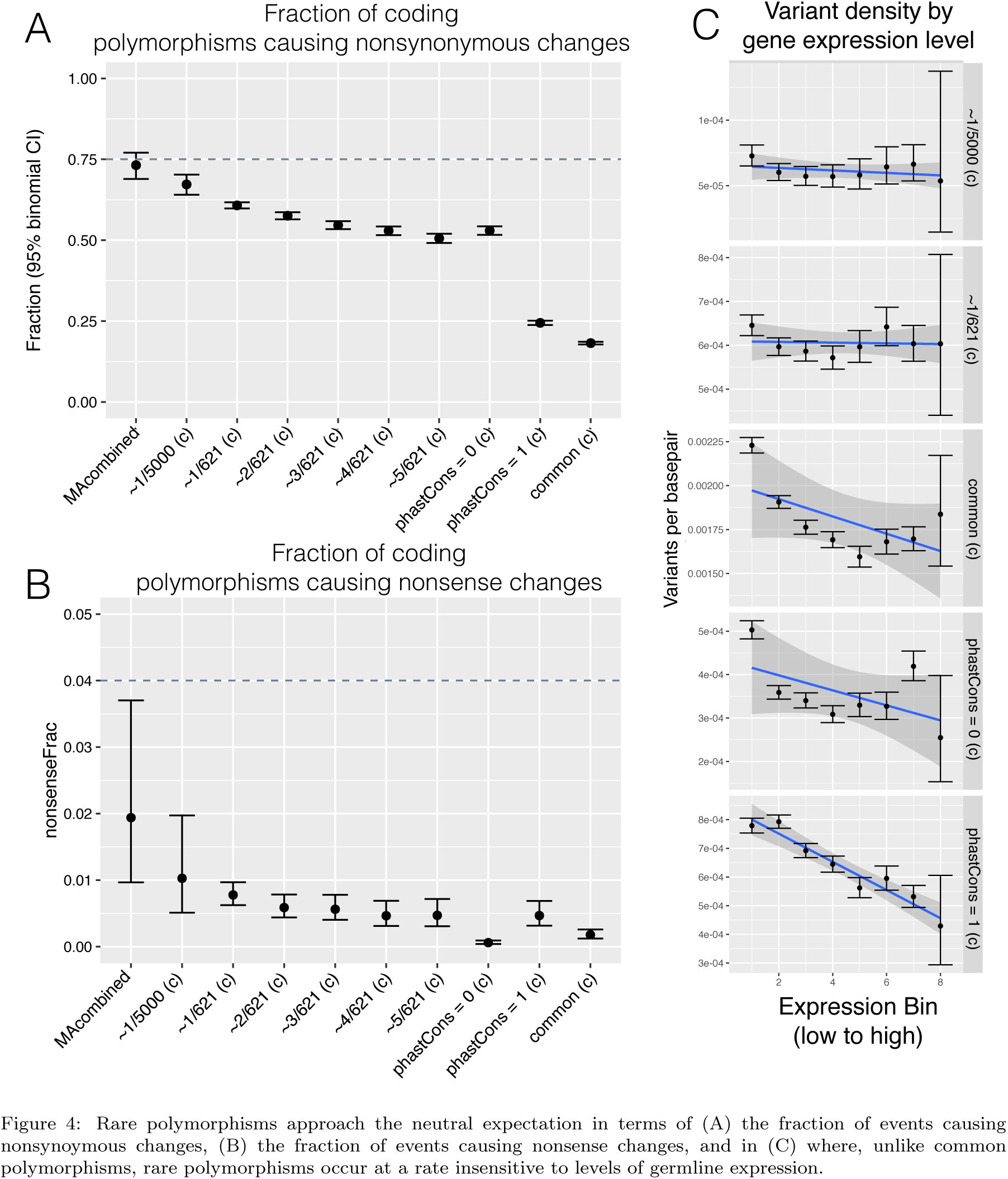
Rare polymorphisms approach the neutral expectation in terms of (A) the fraction of events causing nonsynoymous changes, (B) the fraction of events causing nonsense changes, and in (C) where, unlike common polymorphisms, rare polymorphisms occur at a rate insensitive to levels of germline expression.

#### Rare polymorphisms approach the neutral expectation within coding regions

We next sought to confirm that, in contrast to common polymorphisms, the rarest class of polymorphisms approach the neutral expectation for new mutations. To do this we applied the same tests implemented for the MA experiments to polymorphisms occurring in coding regions (Figure 4). Due to coding regions being the regions most susceptible to natural selection, any biases here will allow a gauge of how closely our rare polymorphisms approximate the neutral expectation.

Recalling that the neutral expectation for coding regions is that ~75% of mutations will cause a nonsynonymous change (an expectation reached by MA experiments at 73.2% (CI 68.9-77.1%) nonsynonymous changes). While common polymorphisms in coding regions only consist of 17.9% (CI 17.7-18.2%) nonsynonymous changes, we find that the rarest polymorphisms are significantly higher with 67.2% (CI 64.1-70.3%) of rare variants in coding regions causing a nonsynonymous change (Figure 4A, G test of independence p-value <2.2e-16), although this is still shy of the netral expectation of 75%. We can also look at the fraction of mutations in coding regions that cause nonsense changes, where the neutral expectation is ~4%. In the MA data the fraction of coding changes which cause a nonsense change is 1.94% (CI 0.97-3.70%) nonsense changes, however the event count is too low to detect whether a substantial fraction of nonsense changes are ‘missing’. When looking at the polymorphism dataset, we find that rare polymorphisms in coding regions have 1.03% nonsense changes (CI 0.51-1.97%, not significantly different from MA dataset). This is significantly higher than within common polymorphisms, which consist of 0.06% (CI 0.05-0.09%) nonsense changes (Figure 4B, G test of independence pvalue=1.9e-8).

Lastly, recalling that phastCons scores are a measure of conservation, we can compare the distribution of phastCons scores in the *D. melanogaster* reference genome to the distribution of scores in sites harboring rare polymorphisms. Compared to common polymorphisms, the rarest frequency classes indeed have a distribution of phastCons scores closer to the neutral expectation (Figure S4).

#### Missing deleterious events among both rare polymorphisms and conserved sites

The analyses described above and depicted in Figure 4 illustrate that rare polymorphisms indeed approach the neutral expectation, however there remains a small ‘missing’ fraction of deleterious events within coding regions, presumably because natural selection is efficient enough to remove them even at rare frequencies. Noting that the rarest frequency class consists of ~67.2% nonsynonymous changes, this is ~8/75 = 11% of nonsynonymous mutations that are likely strongly deleterious, as they were unable to reach a frequency of ~1/5000=0.0002. Similarly, the rarest frequency class consists of only ~1% nonsense changes where we expect 4% from neutrality, and thus approximately three-quarters of the nonsense mutations are missing. Interestingly, frequency class is actually a better indicator than conservation status for whether the spectrum of polymorphisms will approach neutrality: while rare polymorphisms consist of ~67.2% nonsynonymous changes, polymorphisms within the least conserved sites (phastCons=0) only reach a nonsynonymous fraction of ~50.1% (Figure 4A, rightmost points) and thus, despite their nonconserved status, are missing about a third of the nonsynonymous mutations expected from neutrality. This trend is similar for nonsense, where the fraction of nonsense changes within coding regions is closer to the neutral expectation for rare polymorphisms than for nonconserved sites (~1.0% vs 0.5% respectively) (Figure 4B, rightmost points, note ‘nonconserved’ is phastCons=0).

One possible explanation could be that these missing nonsynonymous and nonsense mutations could in fact be recessive lethals, and thus potentially underrepresented in our set of resequenced rare polymorphisms. This could occur because the resequenced lines were exclusively inbred strains, and thus recessive lethals either removed via purifying selection during the inbreeding process, or lying within regions known to be enriched for recessive lethals in repulsion [?], and thus masked in the final genotype calls by virtue of appearing in residually heterozygous regions. To test the second possibility we went back to the DGN raw genotype calls and pulled out singletons occurring in a heterozygous state that were also confirmed in the resequencing data. We found that even in this dataset there are only ~1% nonsense changes and ~60% nonsynonymous changes within coding regions (Figure S5), suggesting that balanced recessive lethals are not a substantial fraction of the missing deleterious mutations. This however does not preclude the possibility that the inbreeding process permitted strongly deleterious recessive alleles to be purged out of the genomes [25].

Lastly, we can gain insight into which biological features are most susceptible to the deleterious effects of mutation by performing a GO analysis. Consider that sites susceptible to deleterious events are expected to be over-represented in conserved sites and under-represented among polymorphisms. Indeed, by performing these tests for over- and under-represented terms, we find similar sets of GO terms across all analyses (Table S7). The 5 GO terms which came out in all analyses included: chromatin assembly or disassembly, cytosol, nucleosome, nucleosome assembly, and protein heterodimerization activity. Note that our analysis of over-represented terms within conserved sites gave similar results to previous studies [26]. Overall, implementing this GO analysis on rare polymorphisms permits additional resolution into which organismal processes are critically important.

#### No evidence for mutagenic effects of transcription

There is evidence in some organisms for transcription-associated mutagenesis [36,37]. This is an effect which appears to be clade-specific, and which may be mediated by the extended changes in DNA strand conformation that occurs during transcription, and/or conflicts that occur between transcription machinery and the machinery of other cellular processes. In some organisms, such as humans, transcription-coupled repair can correct lesions incurred on the transcribed strand, although this repair process can, in and of itself, cause other biases in the mutation spectrum. Trancription-coupled repair is thought to be absent from *D. melanogaster*, and thus mutational patterns we find at highly transcribed genes should reflect the mutation spectrum of the transcription process itself rather than any associated repair activities.

We can test whether transcription is mutagenic in D. melanogaster using our data, by measuring the density of polymorphisms within genes that are expressed in the germline. We would suspect that for common polymorphisms there would be a negative correlation between their density within a gene and expression level, reflecting natural selection purging deleterious mutations from important genes. If our rare polymorphisms indeed capture the neutral expectation then we should find this negative correlation to disappear, and in the case of transcription being mutagenic we should find this correlation to turn positive. To test this we downloaded the publicly available expression data generated from the *D. melanogaster* germline by the modEncode project [38], and measured the density of polymorphisms within genes that are binned by expression level (bin levels 1-8 for low-to-high expression, see Methods). We find that common polymorphisms indeed display a negative correlation between their density and expression level within genes, and this correlation disappears for the rare frequency classes of polymorphisms (Figure 4C). This result confirms again that our rare polymorphism data approach the neutral expectation, and suggests that transcription may have no mutagenic effect in *D. melanogaster*.

### Six relative mutation rates, and their dependency on neighbor context among both rare polymorphisms and evolved four-fold synonymous sites

#### Relative mutation rates within rare polymorphisms have spectra similar to MA data

We have calculated the relative rates of the six mutation types across frequency classes (Figure 5A, coding regions only). We find that while common polymorphisms have a spectra significantly different from the mutations that occurred during MA experiments (Fig. 5 *χ*^2^-test pvalue < 2.2e-16), the rarest polymorphisms have relative mutation rates which approach the MA spectra (Fig. 5A, *χ*^2^-test comparison with MA gives p-values =0.0003 and =0.06 for rare polymorphisms at frequencies ~1/5000 and ~1/621 respectively). Differences between the rare polymorphisms at frequency ~1/5000 and MA data are driven by the *C* → *T*/*G* → *A* mutation type (*χ*^2^-tests without this mutation class give corrected p-values>0.01).

**Figure 5:**
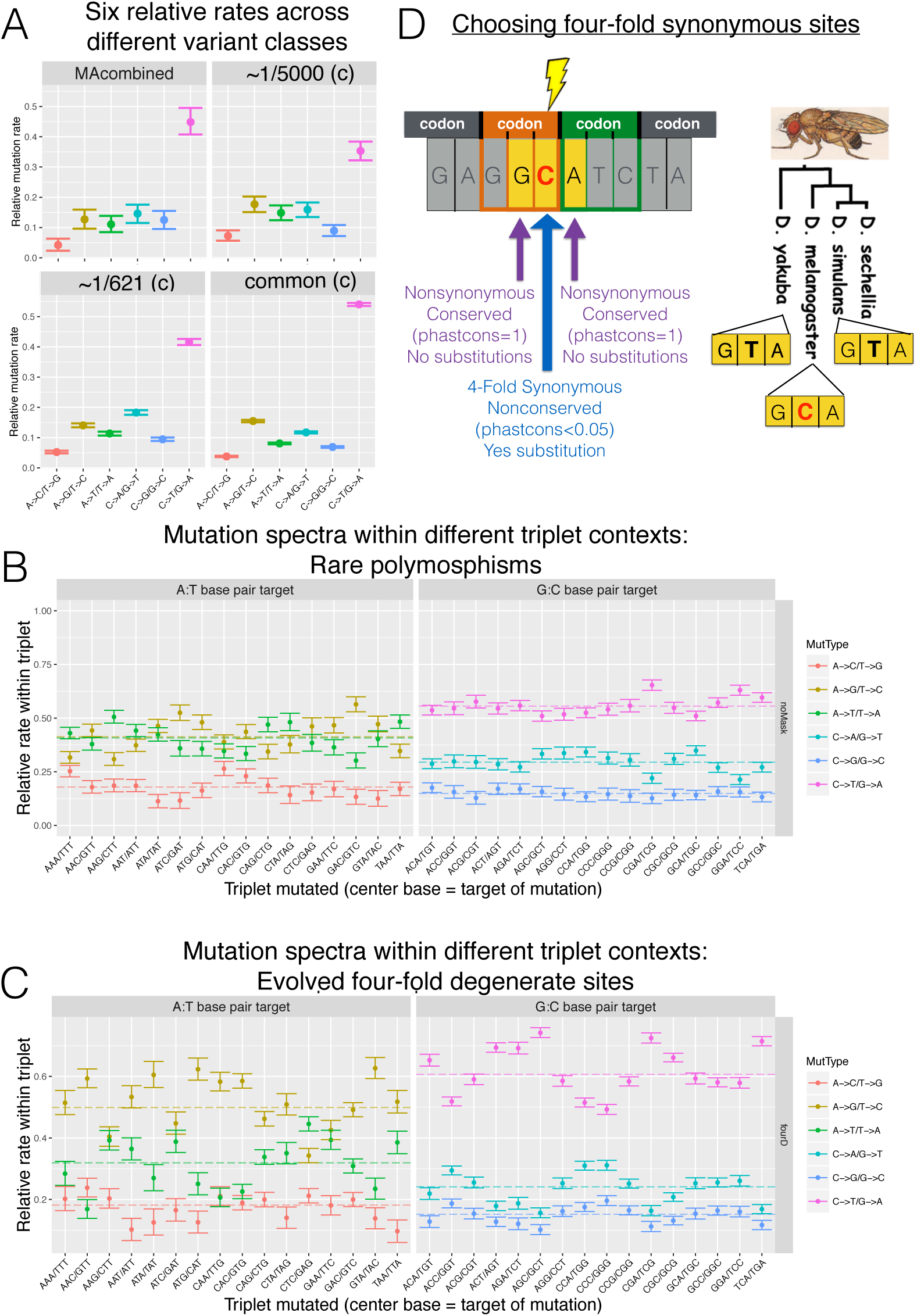
Six relative rates. (A) Six relative rates within MA, rare polymorphisms, and common polymorphisms. (B) Six relative rates within singletons calculated across different triplet contexts, and (C) Six relative rates within substitutions at four-fold synonymous sites, calculated across different triplet contexts. Note the six relative rates within panel (C) are significantly closer to the six relative rates within panel (B) than is expected by chance (p<0.001). (D) Schematic of how four-fold synonymous sites were chosen: the center base of the triplet acquired a substitution on the *D.melanogaster* branch and is conserved in the rest of the *Drosophila* tree, and the outer bases of the triplet is conserved across the entire *Drosophila* tree. Also note, the six relative rates within (C) are significantly closer to the six relative rates within (B) than is expected by chance (p<0.001), indicating that mutational patterns within rare polymorphisms have predictive power for evolution at synonymous sites.

#### Relative mutation rates vary with neighbor-context

Recall that, using the MA dataset of 2, 141 mutations, we were able to detect significant heterogeneity in the mutation spectrum across triplet contexts, however we were unable to detect whether particular triplets were driving the variation (Figure 2E). Now, with our dataset of ~ 70, 000 rare polymorphisms, we can again ask whether the mutational spectrum is dependent on neighbor context.

To this end, we again collapsed all possible triplets into their forward/reverse sequence, and then quantified within each triplet the relative rates of the three mutation types that can occur at the center base pair of the triplet (e.g. CAG/CTG triplet can have A→T/T→A, A→C/T→G, or A→G/T→C mutations). We tested for heterogeneity in the mutation spectrum using rare polymorphisms, and indeed found a quite significant effect of triplet context (Figure 5B, G test, p-value < 2.2e-16 for both GC and AT base pairs). Additionally, we can detect triplet specific effects, where we find 6/16 triplets centered at G:C basepairs to have significant effects, and 14/16 triplets centered at A:T basepairs to have significant effects (G tests, corrected p values < 0.01, Table S8).

#### Neighbor-dependent relative mutation rates predict evolution at four-fold synonymous sites

We wished to test whether our measured context-dependent effects had any predictive power during the course of *Drosophila* evolution. In particular, we were curious whether evolution at four-fold synonymous sites could be predicted, as these sites are known to have particular biases in codon-usage, however the contribution of mutation to these patterns has yet to be fully elucidated [39,40]. This was a particularly intriguing test due to the fact that the neighbors on either side of four-fold sites are, by definition, nonsynonymous, (see schematic in Figure 5D, left), and thus potentially provide a long-term conserved triplet context that could affect evolutionary patterns at the center base pair.

To test this, we identified four-fold synonymous sites within the *D. melanogaster* genome [6] that we could use to measure triplet-context dependent evolution, or in other words nonconserved four-fold sites with conserved neighbors. As depicted in Figure 5D, we required the neighbors of the four-fold synonymous site to have a phastCons score of 1 as well as the same base identity across *D. melanogaster*, *D. simulans* and the outgroup *D. yakuba*. Additionally, we required that the four-fold synonymous site itself to be relatively nonconserved with a phastCons score ≤0.05, and also required that the site had a substitution occur in the *D. melanogaster* branch while simultaneously having no substitution occur on the *D. simulans* or *D. yakuba* branches (i.e. a diallelic site where *D.sim*. and *D.yak*. have the same allele, and *D.mel* has a different allele) (see Methods for more detail).

Using these data we then measured the context-dependent effects similarly to before, where within each triplet we quantified the relative rates of the three substitution types that could occur at that triplet (Figure 5C). We found significant heterogeneity across triplet contexts in the spectra of substitution types in *D. melanogaster* (G test p-value < 2.2e-16 for both A:T and G:C base pairs), as well as significant effects of 10/16 and 12/16 triplets centered at A:T and G:C basepairs respectively (G tests, corrected p values < 0.01, Table S9).

We then wished to test whether the patterns of substitutions were significantly more similar to the patterns of rare polymorphisms than expected by chance. To achieve this, we conducted a permutation test as follows: 1) the total G-value was found by summing G-values for each triplet, where the spectrum of mutation types for rare polymorphisms was the expectation and the spectrum for substitutions was the observed, and then 2) the triplet labels of the substitutions were permuted and the total G value was re-calculated, and 3) this permuted G-value was obtained for 1000 different randomizations. The total G value of the original observed substitution data fell below the zero-percentile of the distribution of G-values for the permuted data. This confirms that mutation, as predicted by the spectrum of rare polymorphisms, does have a significant (*p* < 0.001) impact on the evolution of codon usage at four-fold synonymous sites.

### Equilibrium GC content impacted by neighbor context, but not recombination

#### GC equilibrium as a function of neighboring GC content

It has been observed before that GC-rich regions tend to favor nucleotide changes towards G:C base pairs among common polymorphisms [9], however it is unclear whether this pattern is driven by selective or mutational forces. Thus, we next sought to test for context dependent effects on the mutationally-driven GC content of the genome. We chose to look at GC equilibrium at a site as a function of the GC content of the neighboring base pairs on either side of that site. To this end, we collapse triplet contexts to both strand-indifferent (i.e. an A:T neighbor base pair is the same as a T:A neighbor base pair) and site-indifferent (i.e. the center base can be A, T, C, or G) contexts, such that there are only six contexts total (i.e. see legend of Figure 6A). Note, the only characteristics thus distinguishing these six neighbor contexts is the GC content of the neighboring bases, and how the neighbor base pairs are oriented towards each other. Using these six simple contexts we can calculate the GC equilibrium at the center site using the MA combined dataset and the rare polymorphism dataset. We find a positive correlation between GC equilibrium and the GC content of the neighboring base pairs (Figure 6A), as well as a positive correlation between GC equilibrium and the GC content of the center base pair in the reference genome (Figure S6). These correlations are not significant for the MA combined dataset (p = 0.11 and p = 0.15 respectively), but does meet the significance threshold for the rare polymorphism dataset (p = 0.02 and p = 0.01 respectively). This result suggests two equally interesting possibilities - either mutational forces are contributing to GC-biased nucleotide changes within GC-rich regions or, probably less likely, selection is driving GC-bias in GC-rich regions even within the rarest polymorphism class.

**Figure 6:**
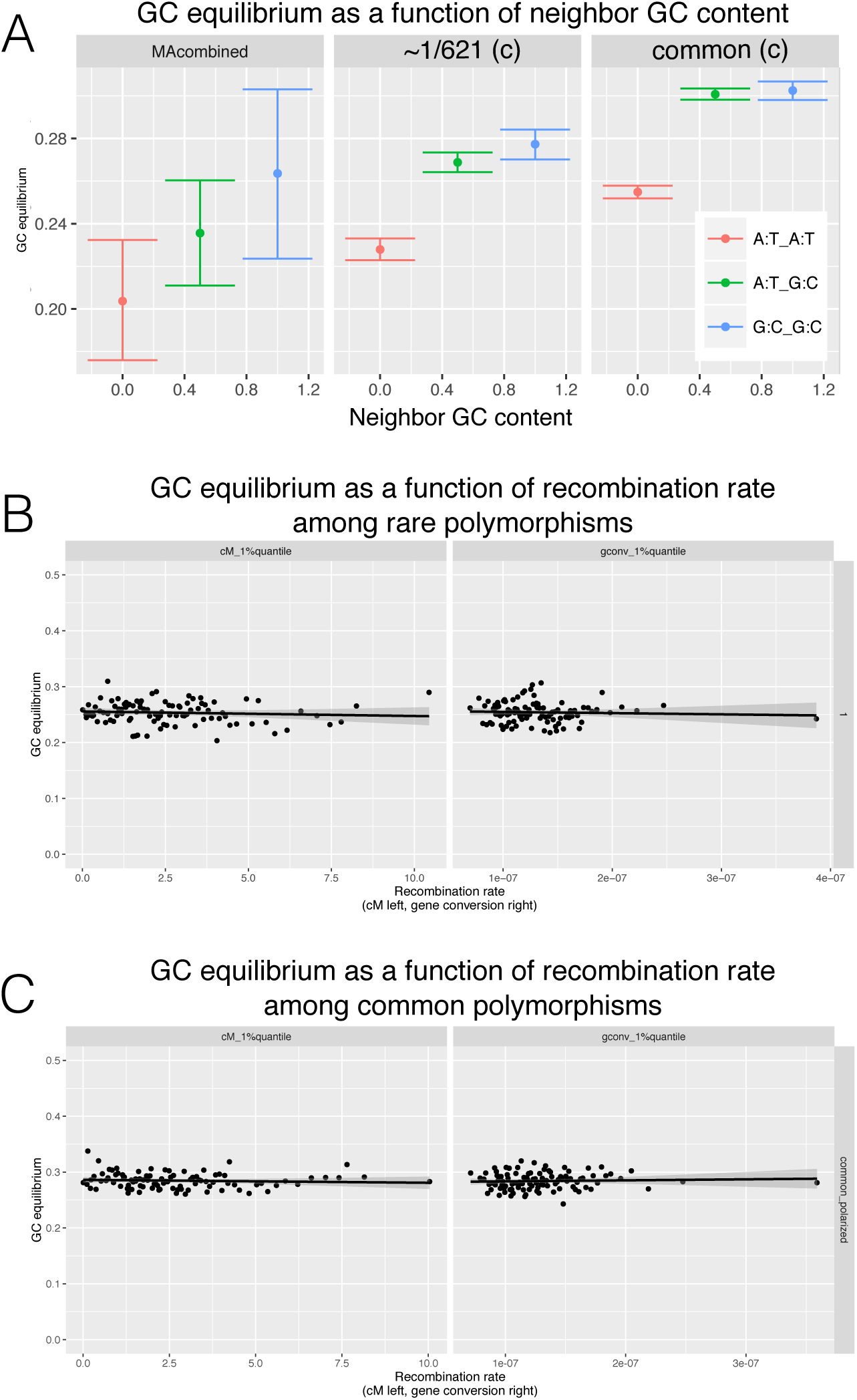
GC equilibrium. (A)GC equilibrium as a function of the GC content of neighboring bases, within MA, singletons, and common polymorphisms (after repetitive regions masked in reference genome). (B) GC equilibrium within singletons and as a function of the recombination rate, and (B) GC equilibrium within common polymorphisms and as a function of the recombination rate.

#### Recombination rate does not affect GC equilibrium

In some organisms it has been found that recombination promotes mutation [41]. It can be difficult to test for whether recombination is mutagenic due to the confounding effect of selection. Natural selection is more efficient in regions of higher recombination and consequently can cause a positive correlation between diversity levels and rates of recombination [42] - the same signature we’d expect to find if recombination is mutagenic. However, we can employ the GC equilibrium metric to test for whether recombination affects the spectrum of mutation types. If, for example as has been found in humans [41], recombination inflates the rate of *C* → *T* transitions relative to non-recombining regions, we would then expect GC equilibrium to decrease with increasing recombination rate. To measure the relationship between recombination and GC equilibrium, we downloaded publicly available genome-wide estimates of both crossover and gene conversion rates [43], and estimated GC equilibrium as a function of these recombination rates using different frequency classes of polymorphisms (Methods). We find no correlation of GC equilibrium with either crossover or gene conversion rates (Figure 6B-C), thus suggesting that recombination does not alter the spectrum of new mutations.

### Multinucleotide mutations comprise ~4-10% of rare polymorphisms, significantly more is than expected by chance

We last sought to test for evidence that singletons cluster together within each strain significantly more than would be expected if all events were independent, indicating multinucleotide mutation events. The first measure we used was the relative proportions of the different type of multinucleotide mutations, which can be seen in Figure 7A and Table 5. The nearest neighbor distance was calculated for every singleton (i.e. distance to the closest neighboring singleton within the same strain), and the expectation was calculated by permuting the strain IDs across all singletons and re-calculating the nearest-neighbor distances for each sample’s singletons, and then taking the average of 500 permutations. As can be seen in Figure 7A and Table 5, 4% of singletons occur in clusters within 1-5bp, where the expectation is only 0.02%. Note that this dramatic enrichment of multinucleotide mutations is robust to a number of strategies for calculating the expected distribution (Figure S7). Among the singletons occurring within 1-5bp of each other, about a quarter of them are ‘duples’, a pair of singletons directly next to each other, and the rest are singletons which occur up to 5bp away from another singleton in the same strain (Table 5). Interestingly, there are even significantly more singletons clustering in the 1-10kb range than is expected by chance, suggesting regional increases in mutation rate may be occurring as well.

**Figure 7:**
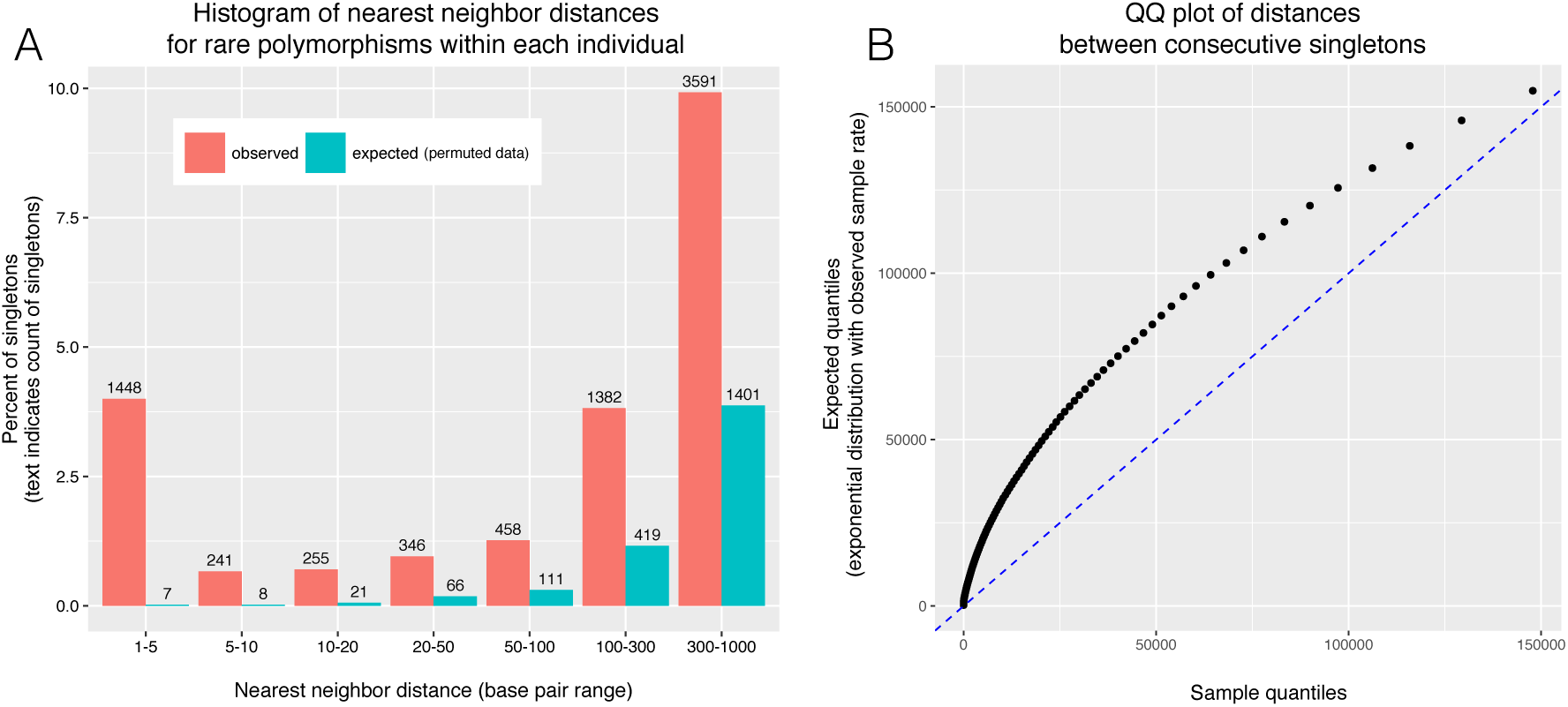
Multinucleotide mutations occur more often than is expected by chance. (A) Histogram of nearest neighbor distance, where every singleton was assigned the distance which was the shorter of the two distances on either side (within a given individual). The expectation is taken from the average of 500 permutations of sample IDs. (B) Quantile-quantile plot of distances between consecutive singletons (on both sides of singletons, within an individual). The expectation is taken from an exponential distribution with a rate equal to the rate within the observed data.

**Table 5:**
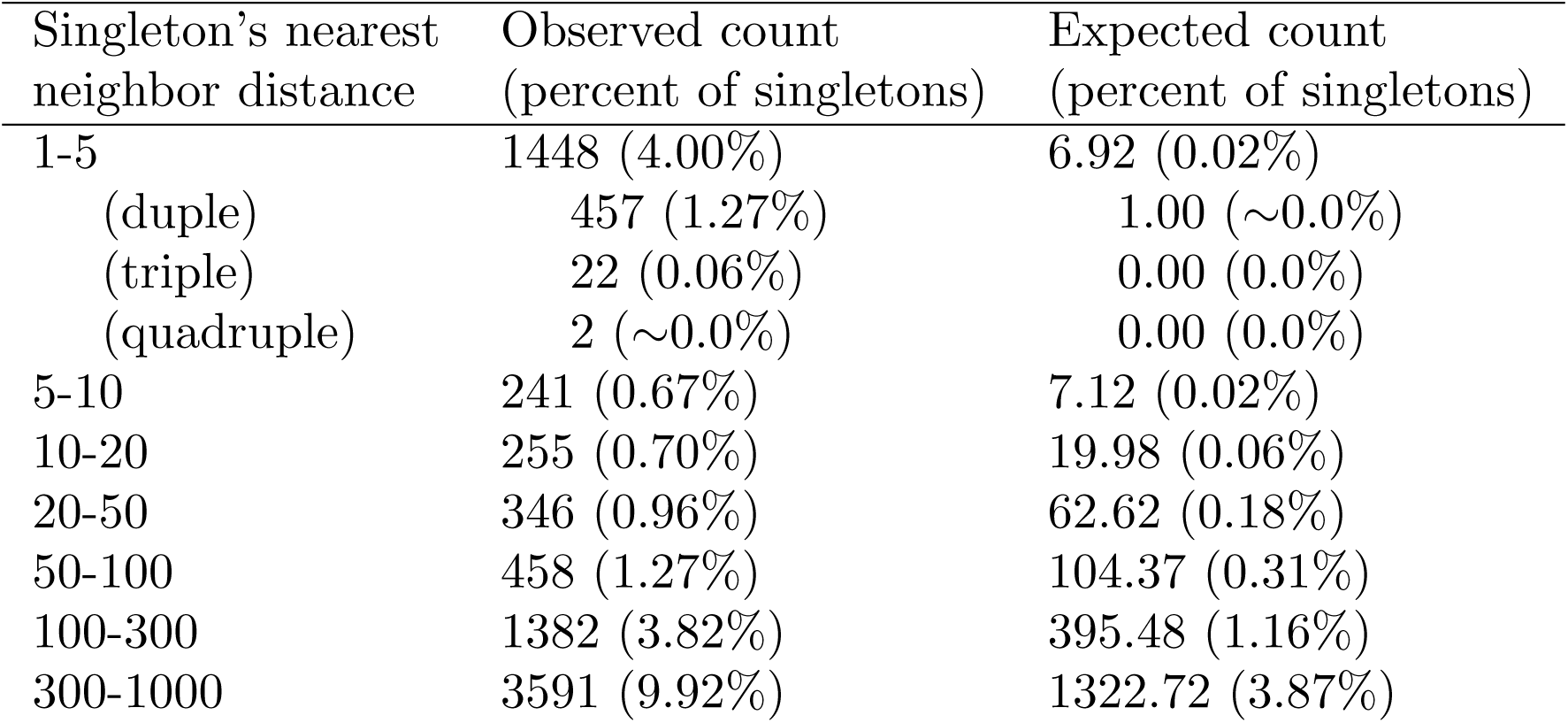
Count of multinucleotide events within singletons (freq~1/621). The expectation was found by permuting strain ID and recalculating the number of events, and taking the average of 500 repetitions of this procedure.

This skew towards shorter distances can also be seen by considering that, if mutations occurred independently, then we expect the the distances between consecutive singletons (within a given individual) to be exponentially distributed. To test this, the quantiles of the distances within the sample data were plotted against the quantiles of an exponential distribution with the a rate equal to the average singleton rate across all strains. As can be seen in Figure 7B, the observed sample data have a distribution that is skewed towards smaller distances.

## Discussion

Mutations provide the raw material of evolution, and thus fundamental to our ability to study genome evolution is the need to have precise measurements of mutational rates and patterns. To this end, we have united multiple strategies, both experimental and computational, to generate the largest and highest quality dataset of *de novo* and rare polymorphisms yet available in *Drosophila*.

### Experimental approach (MA studies)

With respect to the experimental approach, we contribute a novel dataset of 325 *de novo* point mutations generated during a heterozygous mutation accumulation (MA) experiment, and also contribute the first meta-analysis to be done on all sequenced *Drosophila* MA lines. In our metaanalysis, we find that the spectrum of mutation types is remarkably similar across experiments, while the single base pair mutation rate is significantly different. At first glance this seems a surprising result, however upon closer consideration these observations may not be so incongruous.

The published studies which reported lower mutation rates tended to use homozygous accumulation, older technology, and longer generation times. Anywhere from one to all of these factors may be driving the difference in reported rates, but there is no substantial evidence that these particular factors would drive a difference in mutation spectra. For example, if the difference in mutation rates is driven by older vs newer technologies (such that older studies had lower rates of mutation detection), as long as the detection rates of different technologies do not vary by mutation type, then the mutation spectrum should not vary either. Alternatively, if the difference in base pair mutation rates is mediated by the heterozygous vs homozygous experimental design (which, respectively, do and do not allow recessive lethals), there is nonetheless little data supporting the idea that recessive mutations would have a different spectra of mutation types [44,45]. There is also recent evidence from Sharp et. al. (2016) [24] suggesting that the single base pair mutation rate and spectra are robust to experimental design. In their work, the MA experiment was done in different genetic backgrounds (wildtype and deleterious), and they found a difference in the fitness decline over time that was mediated by a difference in the indel mutation rate, not the single base pair mutation rate or spectra. Interestingly, it has also been found in yeast that that the single base pair mutation rate is consistent across MA strains and experiments but the indel mutation rate is not [46]. Overall, we think the significantly different total mutation rates across *Drosophila* MA strains do not reflect a fundamentally different single base pair mutation process across strains, but rather a difference in experimental design and technology.

These observations, in combination with our findings that the single base pair mutation spectra in all experiments fit the neutral expectation, validates the MA approaches for characterizing the spectra of new single base pair mutations, and further validates combining the data across the five experiments and 158 lines into one large dataset. We have made the entire meta-dataset of 3,191 point mutations available for download, along with the curated set of 2,141 point mutations which reside on the major autosomes, outside of repetitive regions, and within non-mutator lines. Using the dataset of 2,141 truly new mutations we have made the highest precision estimates yet available for the spectrum of new mutations in *Drosophila*.

### Computational approach (assaying rare polymorphisms)

We have united disparate genomic resources in the *D. melanogaster* community to generate the first massive set (~70,000) set of high quality, fully-validated, rare polymorphisms, with which we precisely measure mutational patterns across the genome.

The identification of rare polymorphisms in natural populations allows us to circumvent the laborious MA experiment and create a dataset of mutations which, due to being rare, are relatively unaffected by the filter of selection. The challenge to this strategy, however, is distinguishing rare polymorphisms from sequencing and alignment artifacts. To address this challenge, we leveraged the availability of resequence data for 29 strains of *Drosophila melangaster* in order to fully validate our entire polymorphism dataset. We show that the rate of validation of segregating polymorphisms actually decreases with their observed frequency, a result which emphasizes the importance of using resequence data to validate even high coverage genomes. Furthermore, by investigating which genotype calls were not confirmed during resequencing, we find that artifactual calls are more likely to occur in low complexity regions and near sites enriched for segregating indels.

Our finding that rare variants are conflated with artifactual genotype calls at a high rate, even when called in high-coverage genomes and with severe filtering on quality scores, should be of broad interest to the genomics community because the majority of genetic variants segregating in natural populations are rare. For example, many widely-used statistical tests that rely on the site frequency spectrum [47–50] are sensitive to erroneous rare variant calls [19,51–54]. Additionally, there is an ever-expanding catalogue of disease alleles that are found to be private to individuals or families [55–57], and thus erroneous calls can potentially affect patient diagnosis and treatment. Our work reaffirms that artifactual variant calls disproportionately affect rare variants, and that it would be best to incorporate resequencing into any study which analyzes them.

In order to detect whether our dataset provides a valid method for the study of mutational patterns, we looked at whether rare polymorphisms approach the neutral expectation within coding regions. We expect random mutations in coding regions to cause nonsynonymous changes ~75% of the time. While common genetic variants consist of ~20-25% nonsynonymous changes (reflecting the filter of selection), and MA experiments consist of ~75% nonsynonymous changes (reflecting the neutral expectation), we find that our dataset of rare polymorphisms reaches ~68% nonsynonymous changes. This dataset is thus significantly closer to the neutral expectation as compared to genetic variants segregating at higher frequency, confirming that rare polymorphisms provide a reasonable approach for studying mutational patterns.

With our massive set of rare polymorphisms, we detect significant fine-scale heterogeneity in the mutation spectrum across different sequence contexts (‘triplets’). Context-dependency of mutation has been detected in other organisms, including humans [17,24]. Our novel contribution here is, in addition to the highest precision estimate of context-dependency yet available in *Drosophila*, a demonstration that our detected mutational patterns are relevant to the course of evolution within coding regions. The context-dependent rates of mutations, as measured from rare polymorphisms, predict the spectra of substitutions which occurred at four-fold synonymous sites in the *D. melanogaster* phylogenetic branch. This shows that, in addition to forces like selection for translational efficiency [39,58] or biased gene conversion [59,60], the mutation process itself is contributing to biased codon usage patterns at synonymous sites.

Genome-wide GC content in *Drosophila* is ~43%. Using both MA data and rare polymorphisms, we establish that mutational processes by themselves are expected to drive the genome GC content to ~25%. The pattern of mutational GC equilibrium being significantly lower than actual genome GC content is common across the tree of life [10], and presents a question of which forces drive the genome GC content to such high values in general, and in *Drosophila* specifically. Although in many species weak selection and/or biased gene conversion were implicated in the evolution of high GC content, this explanation is unlikely to be valid in *Drosophila*. This is because the common polymorphisms do not display a substantial bias towards higher GC values (~28%), and neither does this bias increase with recombination rates. Both of these patterns would be expected under the models of weak selection or biased gene conversion. It is likely instead that the high GC content of the *Drosophila* genome reflects its high functional density and the elevated GC content of those functional sequences. Consistent with that model, the parts of the genome that are expected to have lower functional density do have substantially lower GC content. For example, the average GC content of short introns is ~32% [61], indicating that the even the best neutral-standard available for Drosophila may still have some slight constraint.

We observe another interesting relationship between genome GC content and GC equilibrium - a correlation between the mutational GC equilibrium and the local genome GC content, such that mutational processes drive GC content to lower levels in already GC-poor regions. It has been observed before that common polymorphisms in intergenic regions display this same pattern of a lower GC equilibrium in GC-poor regions [9,61]. It was thought that such a pattern is largely driven by selective forces. However, our dataset of *de novo* MA mutations and also rare polymorphisms is large enough to show that the pattern persists even among genetic variants that have little-to-no filtering from natural selection (or biased gene conversion). Thus, while mutational processes drive genome GC content down and selective forces drive genome GC content up, we find that mutation is most strongly biased to lowering GC content in already GC-poor regions.

Lastly, we were able to show that multinucleotide mutations occur significantly more than is expected by chance, and we have precisely quantified the relative rates of each type of multinucleotide event. In *Drosophila* MA studies which used a set of ~ 1000 *de novo* mutations [22,24], it has been found that about ~3-4% of single base pair events occur in clusters of size ≤50bp, coinciding closely with our finding that ~4% of events occur in clusters of ≤5bp and another ~2% in 6-50bp. Similar rates of small multinucleotide clusters have been found in other organisms [62–64]. With our larger dataset we can take the analysis a step further and find that as many as ~10% of single base pair mutations occur in clusters of size ≤ 1kb (where the expectation is only ~4%). This result is consistent with some recent work done in humans [14,62] which also suggests that such regional increases in mutation rate may be a common occurrence in the genome.

In combination, the MA approach and the rare polymorphism approach have provided complementary methods for studying the spectrum of new mutations, enabling precise estimate of both total mutation rates and subtle mutational biases. We hope, with an ever-growing catalog of deep sequence data from natural populations being made available to the scientific community, that researchers will take advantage of the opportunity to apply the methods described here to studying mutational patterns in other organisms.

## Materials and Methods

### Mutation accumulation

Two strains are used for the experiment: DGRP strain RAL-765 (wildtype genotype 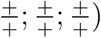 which has red eyes, and a white-eyed laboratory strain 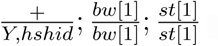 generated by Mark Siegal at New York University, hereafter referred to as hshid (for the ‘heat shock head involution defective’ genotype). The hshid strain allows the easy acquisition of virgins, because all males die upon one-hour heath shock during the larval stage. Using these two strains, the MA chromosomes (i.e. DGRP) are always passed through red-eyed males that are heterozygous with the hshid chromosomes, allowing the MA chromosomes to avoid recombination (because males lack it) and to never be subjected to heat-shock (because only hshid chromosomes are heat-shocked). The crosses were as follows: A single male fruit fly from DGRP strain RAL-765 was crossed with 6 virgin hshid females, after which a single red-eyed male progeny (i.e. heterozgous for both bw and st and therefore carrying the founder male’s 2nd and 3rd chromosomes) was taken and crossed to 6 hshid virgin females. From this cross, 50 red-eyed male progeny were then isolated within vials, thus founding 50 MA strains with identical chromosome 2 and 3. Thereafter every generation a single red-eyed male progeny from each strain was crossed to 3 hshid virgin females, and as backup this was replicated using male siblings in 3 additional vials each generation.

### MA sequencing

As is common in MA experiments, some lines became sick and died off as generations passed, such that we were able to sequence 17 lines out of the original 50. From these 17 MA strains, five to fifteen flies were taken (thus heterozygous for the MA chromosome) in generations 36, 37, 49, and 53, and DNA extracted according to a standard protocol [65]. Paired-end barcoded DNA sequencing libraries were prepared using illumina Nextera Sample Preparation Kit (FC-121-1031) and Index Kit (FC-121-1012) and KAPA Biosystems Library Amplification Kit (KK2611), modified for small volumes. In brief: Genomic DNA was diluted to 2.5ng/uL using quantification with Qubit HS Assay Kit (Q32851), and then 1ul of DNA was ‘tagmented’ according to the Nextera protocol in a 2.5ul total volume (55°C, 5 minutes), then each library was amplified using the KAPA amplification kit with Nextera index primers (N7XX and N5XX) in a 7.5ul total volume (98°C for 165sec, 8 cycles of 98°/62°/72° for 15sec/30sec/90sec), and then reconditioned to add Nextera PCR primers (required for illumina sequencing) in a 17ul total volume (95°C for 5min, 4 cycles of 98°/62°/72° for 20sec/20sec/30sec). Libraries were sequenced on a HiSeq 2000 to a depth of 20-25X (see SM for sample coverage plot).

### MA variant calling

In order to call *de novo* mutations in these strains heterozygous for the MA and hshid chromosomes, we sequenced the ancestral DGRP and hshid strains in addition to each MA line, and then accepted genetic variants which were strictly unique to each MA strain (thus hshid variants segregating across the heterozygous MA strains were filtered out). The following pipeline was used to call genetic variants in the MA sequencing data, where default settings for each tool were used unless otherwise specified: Reads were trimmed with TrimGalore! v0.3.7 [http://www.bioinformatics.babraham.ac.uk/projects/trimgalore] (`trim_galore −a CTGTCTCTTATACACATCT −a2 CTGTCTCTTATACACATCT --quality 20 --length 30 --clip_R1 15 --clip_R2 15 --three_prime_clip_R1 3 --three_prime_clip_R2 3 --paired`), then mapped to release 5.57 of the Drosophila reference genome with BWA-MEM v0.7.5a [66]. Next PCR duplicates were removed with PicardTools v1.105 [http://broadinstitute.gith (`MarkDuplicates REMOVE DUPLICATES=true`), and then reads locally rearranged around indels with GATK v3.2.2 [67] (`RealignerTargetCreator` and `IndelRealigner` tools).Variants were called with GATK (`HaplotypeCaller --heterozygosity 0.01`) followed by the GATK recommended conservative filters (`VariantFiltration --filterExpression "QD <2.0 || FS > 60.0 || MQ < 40.0 || MQRankSum < −12.5 || ReadPosRankSum < −8.0"`), and new mutations were considered the variants present in one MA strain and absent from all the rest. Variants were also called with a less conservative pipeline using SAMtools v0.1.19 [68] to generate a mpileup file (`mpileup`) followed by Varscan v2.3.9 [69] to call variants (`mpileup2cns --min-coverage 4 --min-reads2 2 --min-var-freq 0.01 --min-freq-for-hom 0.99 --strand-filter 0 --p-value 1`), and again new mutations were considered the variants present in one MA strain and absent from all the rest. Repetitive regions were then filtered out, including from RepeatMasker [http://www.repeatmasker.org], from a run of TRF [70] on the Drosophila reference, and from a list of annotated transposable elements [71]. The final list of new mutations was considered the intersection of the two variant call sets, in order to ensure that a new mutation was both a high quality variant call (i.e. the conservative GATK set) and also never observed in any other strain (i.e. nonconservative Varscan set). Thirty variants were amplified with PCR and then Sanger sequenced, of which twenty-nine were confirmed. A summary of the mutation counts and generations of mutation accumulation for each strain can be found in SM, along with a summary of the mutations randomly chosen for validation via PCR/Sanger sequencing and their corresponding primers.

### MA data from references [21–24]

Lists of mutations from each experiment were downloaded from each publication and then processed with in-house *perl* and *R* scripts to generate a single VCF of all mutations, which is available as a supplementary file. For the ‘MA combined’ datasets, we filtered out repetitive regions (including from RepeatMasker [http://www.repeatmasker.org], from a run of TRF [70] on the Drosophila reference, and from a list of annotated transposable elements [71]), removed mutator lines (line 19 from Huang et. al., and lines 33-27, 33-45, 33-5, and 33-55 from Schrider et. al.), and subsetted down to the major autosomes 2 and 3 only. For comparisons of the single base pair mutation rates with a poisson exact test we require a time base in order to scale the different experiment counts, and while the number of generations and number of lines were provided within the publications, the information on genome size was incomplete. Thus, given that mutation rate *μ* = *m*/(*n* × *t* × *l*) (where *m*=mutation count, *n*=number of strains, *t*= number of generations, and *l* = number of base pairs), we back-calculated *l*, which can be found in the last column of SM.

### Analyses in R

VCFs for all MA experiments were generated with *perl* scripts and then loaded into R with Bioconductor. The majority of downstream analyses were performed with Bioconductor tools, where the genome object `BSgenome.Dmelanogaster.UCSC.dm3` [?] and the transcript annotation object `TxDb.Dmelanogaster.UCSC.dm3. ensGene` [?] were used as references. The functional impact of variants was annotated using the `predictCoding` and `locateVariants` tools.

### DGN rare variant calling

Sequences of the 623 genomes provided by the Drosophila Genome Nexus [31] were downloaded and converted into VCFs, repeat regions (as described above) and were masked, and singletons, doubletons, etc were identified. Additionally, the DGN indel VCFs were downloaded and sites with segregating indels were masked in all strains for the length of the segregating indel plus five base pairs on either side. Variants were confirmed in a subset of strains that had resequence data available from the DPGP1 project’s Solexa (now illumina) sequencing [www.dpgp.org/dpgp_solexa_r1.0.tar], and from the DGRP [72] project’s 454 sequencing data [ftp://ftp.hgsc.bcm.edu/DGRP/freeze1_July_2010/snp_calls/454/]. Singletons confirmed in this way were further filtered down into a set of extremely rare variants by removing any which appeared in NESCENT [?] data, a project which sequenced X wild-caught flies from X populations in North America. These extremely rare variants (‘singletons-noPool’) were removed from the singleton set, in order to create non-overlapping variant sets.

### Variant annotation and analysis

VCFs for MA and DGN data were loaded into R with Bioconductor. The majority of downstream analyses were performed with Bioconductor tools, where the genome object `BSgenome.Dmelanogaster.UCSC.dm3` [?] and the transcript annotation object `TxDb.Dmelanogaster.UCSC.dm3. ensGene` [?] were used as references. The functional impact of variants was annotated using the `predictCoding` and `locateVariants` tools.

## Supplemental Materials, Figures, and Tables

**Figure SuppFigure-1:**
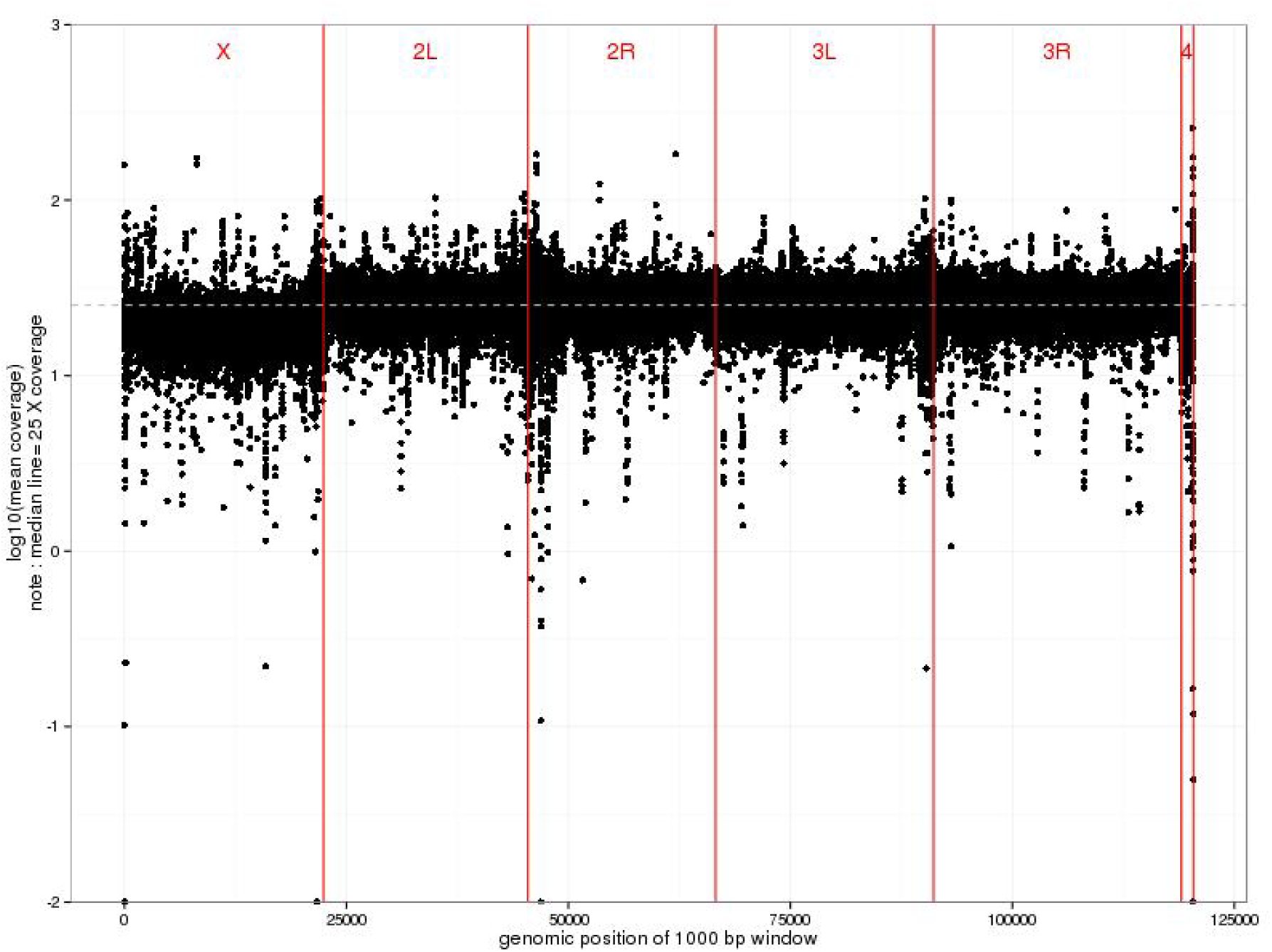
Representative example of the coverage achieved across the genome for a given MA line in this study, here from MA line 33 with a median coverage of 25X.

**Table SuppTable-1:** A summary of the strains sequenced in this study, including the generation at which each strain was sequenced, and the number of mutations identified for each strain.

| Line | Generation<br>sequenced | Total #<br>mutations | chr2L<br>count | chr2R<br>count | chr3L<br>count | chr3R<br>count |
| --- | --- | --- | --- | --- | --- | --- |
| MA_6 | 36 | 17 | 5 | 0 | 5 | 7 |
| MA_9 | 36 | 15 | 3 | 3 | 4 | 5 |
| MA_11 | 36 | 11 | 4 | 2 | 1 | 4 |
| MA_14 | 49 | 32 | 9 | 5 | 7 | 11 |
| MA_16 | 53 | 27 | 7 | 5 | 9 | 6 |
| MA_17 | 37 | 14 | 3 | 4 | 4 | 3 |
| MA_20 | 53 | 21 | 5 | 4 | 5 | 7 |
| MA_21 | 53 | 27 | 3 | 9 | 7 | 8 |
| MA_22 | 36 | 15 | 7 | 0 | 2 | 6 |
| MA_24 | 36 | 19 | 13 | 1 | 3 | 2 |
| MA_27 | 36 | 14 | 3 | 2 | 3 | 6 |
| MA_28 | 36 | 10 | 1 | 6 | 3 | 0 |
| MA_29 | 53 | 21 | 8 | 3 | 6 | 4 |
| MA_30 | 53 | 21 | 2 | 7 | 2 | 10 |
| MA_33 | 53 | 28 | 7 | 1 | 12 | 8 |
| MA_35 | 53 | 15 | 1 | 4 | 4 | 6 |
| MA_50 | 53 | 18 | 5 | 3 | 6 | 4 |

**Table SuppTable-2:**
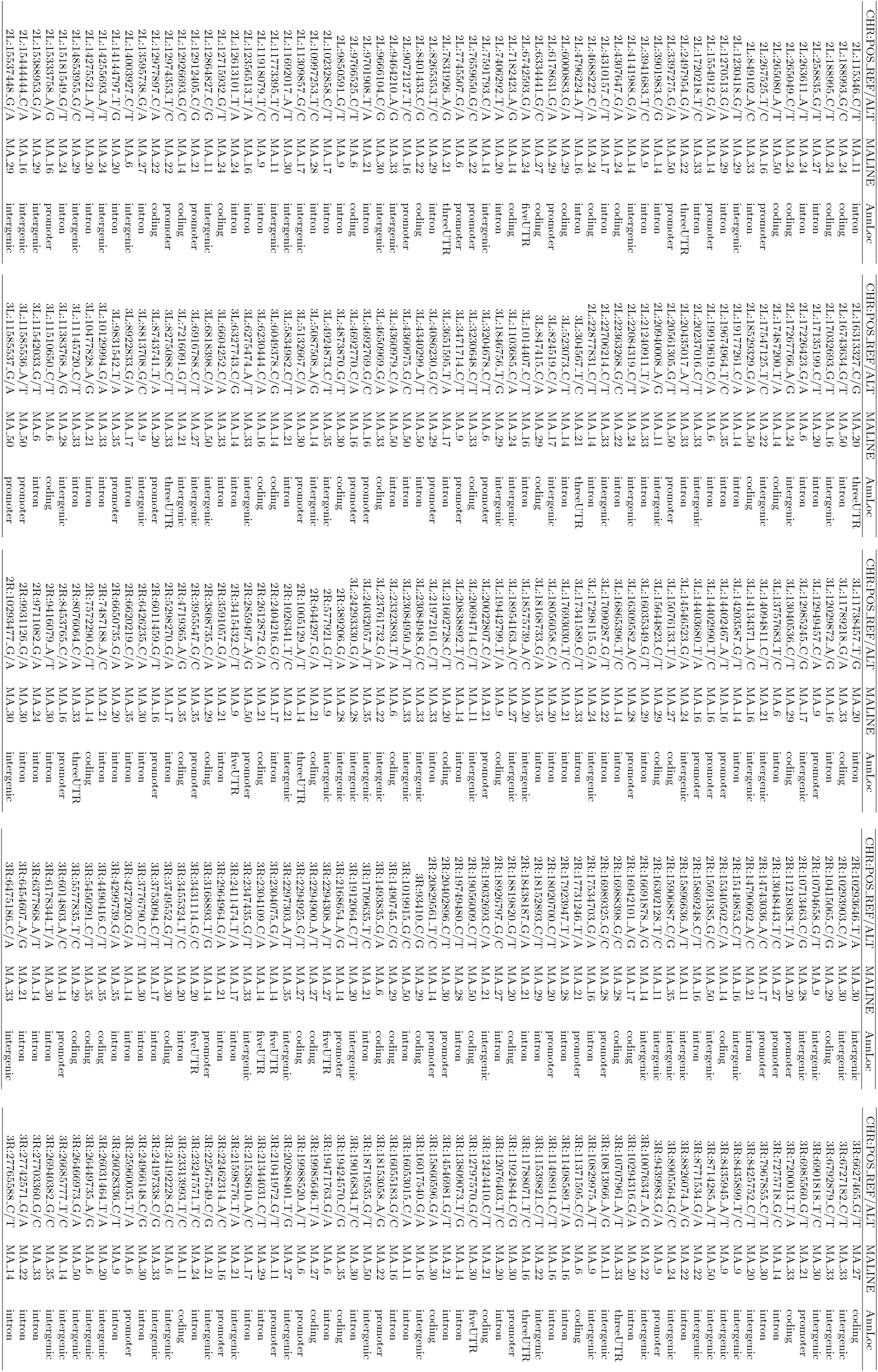
A list of all the *de novo* mutations generated in this study.

**Table SuppTable-3:**
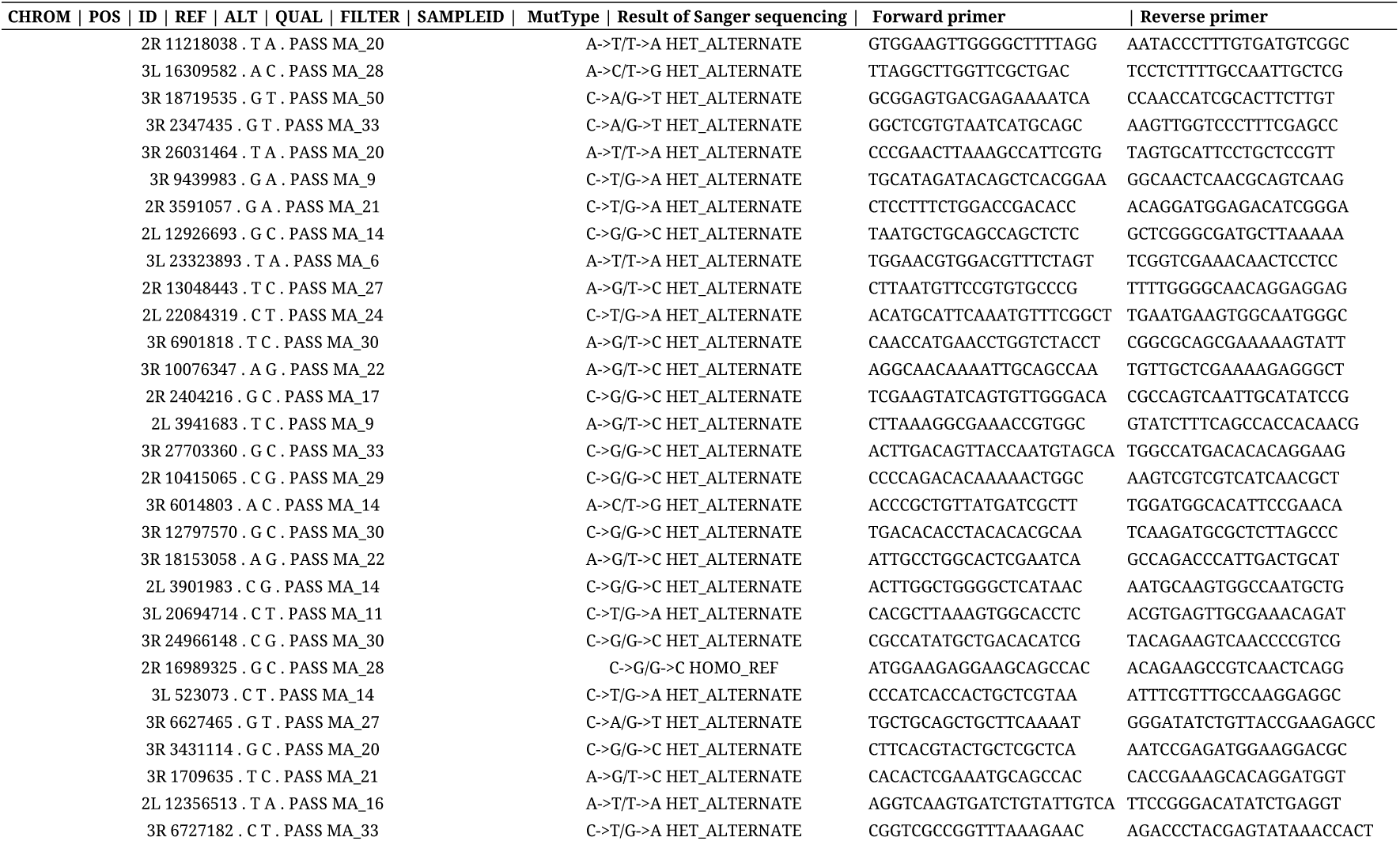
A summary of the 30 mutations chosen randomly from our MA experiment for validation via PCR/Sanger sequencing. This resulted in either a double peak including the reference and alternate allele (’HET_ALTERNATE’) which validates the new mutation, or a single peak matching the reference allele (’HOMO_REF’) which is inconclusive.

**Figure SuppFigure-2:**
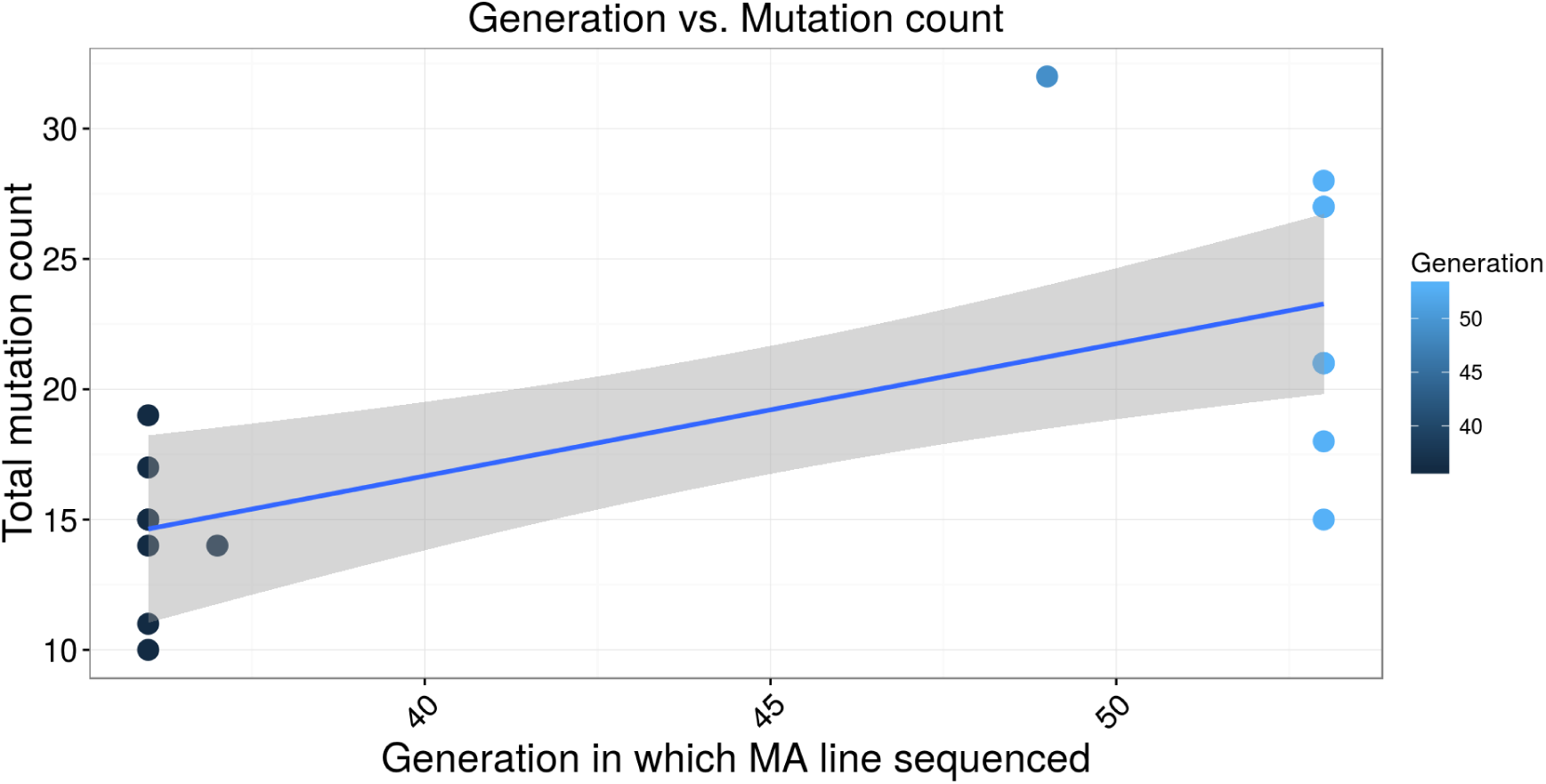
A plot of the total count of mutations on each chromosomal arm for each strain as a function of generation time in which sequenced. There was no significant difference in the rates between the generations, Poisson exact test p-value = 0.757.

**Table SuppTable-4:**
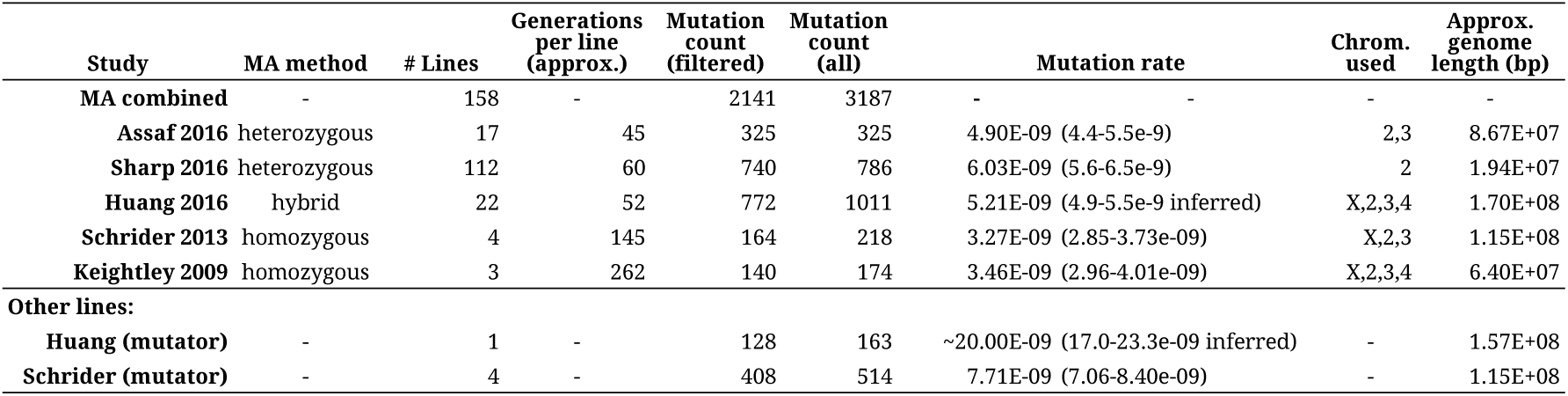
A version of Table 1, with two extra columns indicating the chromosomes used in the experiment, and the back calculated genome length (lengths which were not consistently published within the MA papers). These were used for calculating a time base in the poisson exact test comparing the different single base pair mutation rates across experiments, and were found as follows: given that mutation rate *μ* = *m*/(*n* × *t* × *l*) (where *m*=mutation count, *n*=number of strains, *t*= number of generations, and *l* = number of base pairs), we back-calculated *l*.

**Table SuppTable-5:** A summary of the six relative rates across the five MA studies, as well as a combined estimate. Note this is for the major autosomes 2 and 3 and with repeats masked. The 95% confidence intervals are within the parentheses, and were calculated using 1000 bootstraps of the raw counts.

| Experiment | A → C/T → G | A → G/T → C | A → T/T → A | C → A/G → T | C → G/G → C | C → T/G → A |
| --- | --- | --- | --- | --- | --- | --- |
| MA combined | 0.06 (0.05-0.07) | 0.11 (0.1-0.12) | 0.12 (0.11-0.13) | 0.18 (0.16-0.2) | 0.13 (0.12-0.15) | 0.4 (0.38-0.42) |
| Assaf 2016 | 0.04 (0.02-0.07) | 0.14 (0.1-0.17) | 0.13 (0.1-0.16) | 0.19 (0.15-0.23) | 0.16 (0.12-0.2) | 0.34 (0.29-0.4) |
| Sharp 2016 | 0.06 (0.04-0.07) | 0.11 (0.09-0.13) | 0.12 (0.1-0.14) | 0.18 (0.15-0.21) | 0.1 (0.08-0.12) | 0.44 (0.41-0.48) |
| Huang 2016 | 0.06 (0.04-0.07) | 0.09 (0.07-0.11) | 0.11 (0.09-0.13) | 0.18 (0.15-0.21) | 0.16 (0.13-0.19) | 0.4 (0.37-0.44) |
| Schrider 2013 | 0.08 (0.04-0.12) | 0.15 (0.1-0.2) | 0.14 (0.1-0.2) | 0.19 (0.13-0.25) | 0.1 (0.05-0.14) | 0.34 (0.27-0.41) |
| Keightley 2009 | 0.11 (0.06-0.16) | 0.14 (0.1-0.21) | 0.1 (0.05-0.15) | 0.18 (0.11-0.24) | 0.1 (0.06-0.16) | 0.37 (0.29-0.45) |
| **Other lines:** |  |  |  |  |  |  |
| Huang (mutator) | 0.04 (0.01-0.07) | 0.21 (0.15-0.29) | 0.05 (0.01-0.08) | 0.19 (0.12-0.26) | 0.04 (0.01-0.08) | 0.47 (0.38-0.56) |
| Schrider (mutator) | 0.04 (0.02-0.05) | 0.08 (0.06-0.11) | 0.08 (0.05-0.1) | 0.09 (0.07-0.12) | 0.07 (0.05-0.1) | 0.64 (0.59-0.68) |

**Table SuppTable-6:** The results of G tests for triplet effect on the mutation spectrum, here for the MA combined dataset. From left to right the columns refer to: variant dataset, triplet of interest (forward/reverse), mutation type 1, mutation type 2, mutation type 3, p value for G goodness of fit test (expected is total counts), corrected p value, and whether there is a significant effect of the triplet (p<0.01)

| ALTCOUNT | tripCombo | C->A/G->T | C->G/G->C | C->T/G->A | GtestPvalue | pCORR | pSIG |
| --- | --- | --- | --- | --- | --- | --- | --- |
| MAcombined | ACA/TGT | 18 | 27 | 51 | 4.00E-02 | 6.50E-01 | FALSE |
| MAcombined | ACC/GGT | 14 | 11 | 31 | 9.70E-01 | 1.50E+01 | FALSE |
| MAcombined | ACG/CGT | 11 | 9 | 42 | 1.70E-01 | 2.70E+00 | FALSE |
| MAcombined | ACT/AGT | 20 | 18 | 45 | 7.30E-01 | 1.20E+01 | FALSE |
| MAcombined | AGA/TCT | 26 | 23 | 49 | 3.60E-01 | 5.80E+00 | FALSE |
| MAcombined | AGC/GCT | 29 | 26 | 49 | 1.30E-01 | 2.00E+00 | FALSE |
| MAcombined | AGG/CCT | 17 | 13 | 38 | 9.80E-01 | 1.60E+01 | FALSE |
| MAcombined | CCA/TGG | 41 | 11 | 71 | 6.80E-03 | 1.10E-01 | FALSE |
| MAcombined | CCC/GGG | 19 | 10 | 44 | 5.60E-01 | 9.00E+00 | FALSE |
| MAcombined | CCG/CGG | 23 | 6 | 34 | 5.30E-02 | 8.40E-01 | FALSE |
| MAcombined | CGA/TCG | 17 | 13 | 49 | 5.60E-01 | 9.00E+00 | FALSE |
| MAcombined | CGC/GCG | 15 | 16 | 36 | 5.00E-01 | 8.00E+00 | FALSE |
| MAcombined | GCA/TGC | 40 | 19 | 53 | 5.50E-02 | 8.80E-01 | FALSE |
| MAcombined | GCC/GGC | 22 | 11 | 52 | 4.00E-01 | 6.30E+00 | FALSE |
| MAcombined | GGA/TCC | 15 | 21 | 67 | 2.50E-02 | 4.00E-01 | FALSE |
| MAcombined | TCA/TGA | 29 | 21 | 72 | 8.20E-01 | 1.30E+01 | FALSE |
| MAcombined | AAA/TTT | 29 | 21 | 44 | 1.90E-03 | 3.00E-02 | FALSE |
| MAcombined | AAC/GTT | 7 | 18 | 21 | 6.20E-01 | 9.80E+00 | FALSE |
| MAcombined | AAG/CTT | 10 | 23 | 28 | 6.20E-01 | 9.90E+00 | FALSE |
| MAcombined | AAT/ATT | 7 | 16 | 25 | 2.60E-01 | 4.20E+00 | FALSE |
| MAcombined | ATA/TAT | 11 | 20 | 13 | 3.00E-01 | 4.80E+00 | FALSE |
| MAcombined | ATC/GAT | 6 | 14 | 16 | 8.20E-01 | 1.30E+01 | FALSE |
| MAcombined | ATG/CAT | 3 | 19 | 22 | 3.70E-02 | 6.00E-01 | FALSE |
| MAcombined | CAA/TTG | 13 | 26 | 14 | 9.00E-02 | 1.40E+00 | FALSE |
| MAcombined | CAC/GTG | 8 | 15 | 9 | 3.30E-01 | 5.30E+00 | FALSE |
| MAcombined | CAG/CTG | 8 | 12 | 24 | 1.60E-01 | 2.60E+00 | FALSE |
| MAcombined | CTA/TAG | 7 | 19 | 14 | 5.20E-01 | 8.40E+00 | FALSE |
| MAcombined | CTC/GAG | 6 | 17 | 11 | 4.00E-01 | 6.40E+00 | FALSE |
| MAcombined | GAA/TTC | 12 | 11 | 14 | 2.20E-01 | 3.50E+00 | FALSE |
| MAcombined | GAC/GTC | 6 | 22 | 14 | 1.90E-01 | 3.00E+00 | FALSE |
| MAcombined | GTA/TAC | 6 | 17 | 8 | 1.50E-01 | 2.30E+00 | FALSE |
| MAcombined | TAA/TTA | 14 | 17 | 26 | 3.70E-01 | 6.00E+00 | FALSE |

**Figure SuppFigure-3:**
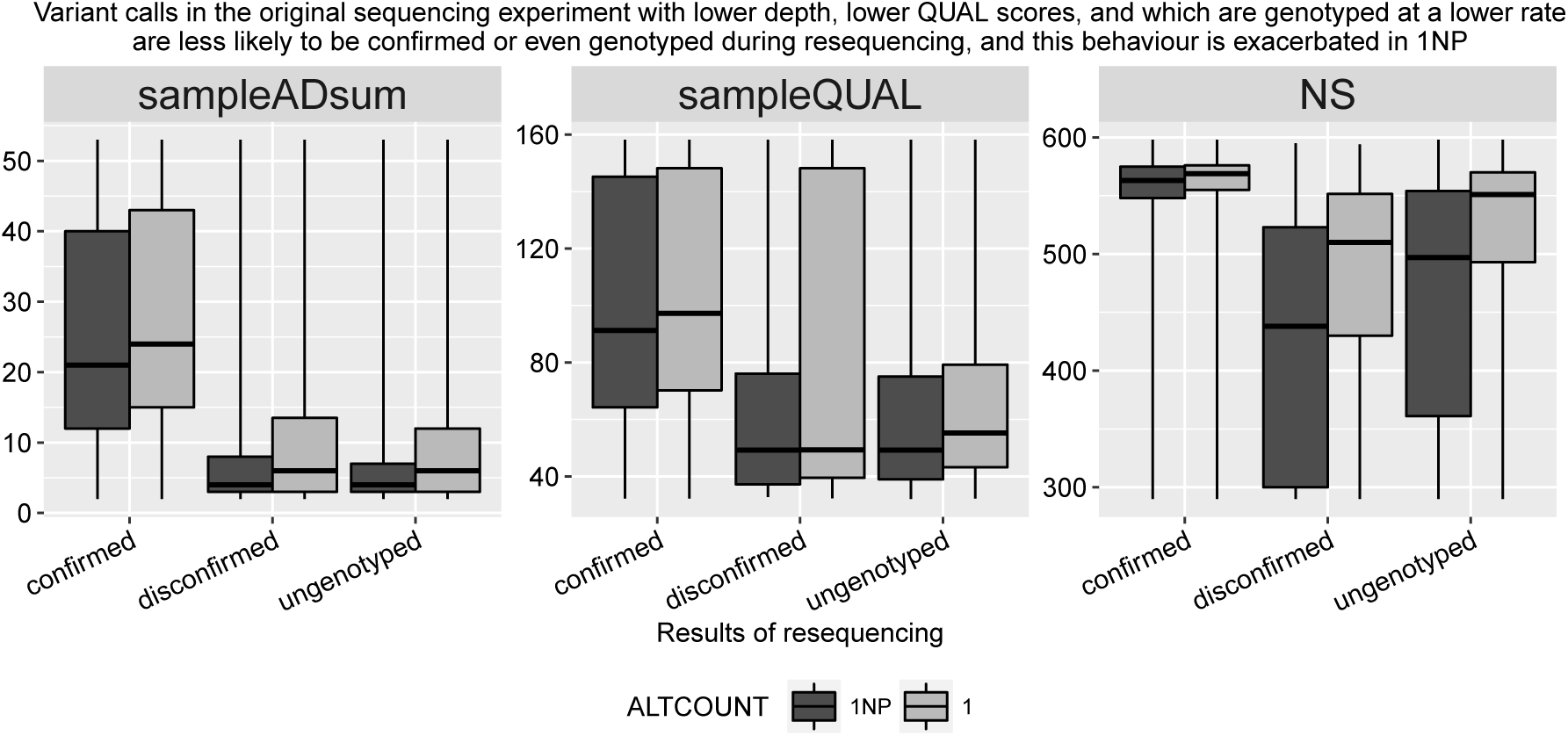
Quantifying the DGN quality metrics of the DGN variants (1NP and 1 refer to variants at frequency ~ 1/5000 and ~ 1/621 respectively) after classifying them by whether in the resequence data the DGN variants were confirmed, disconfirmed, or ungenotyped. Left panel is the total depth at the site in the DGN data, middle is the QUAL score of the genotype call in the DGN data, and right panel is the number of samples with genotype information in the DGN data. DGN variants which were confirmed in the resequence data consistently have higher quality metrics, however there is also overlap in the distribution of DGN quality scores (i.e. DP, QUAL, NS) between those variants which were confirmed, disconfirmed, and ungenotyped in the resequence data.

**Figure SuppFigure-4:**
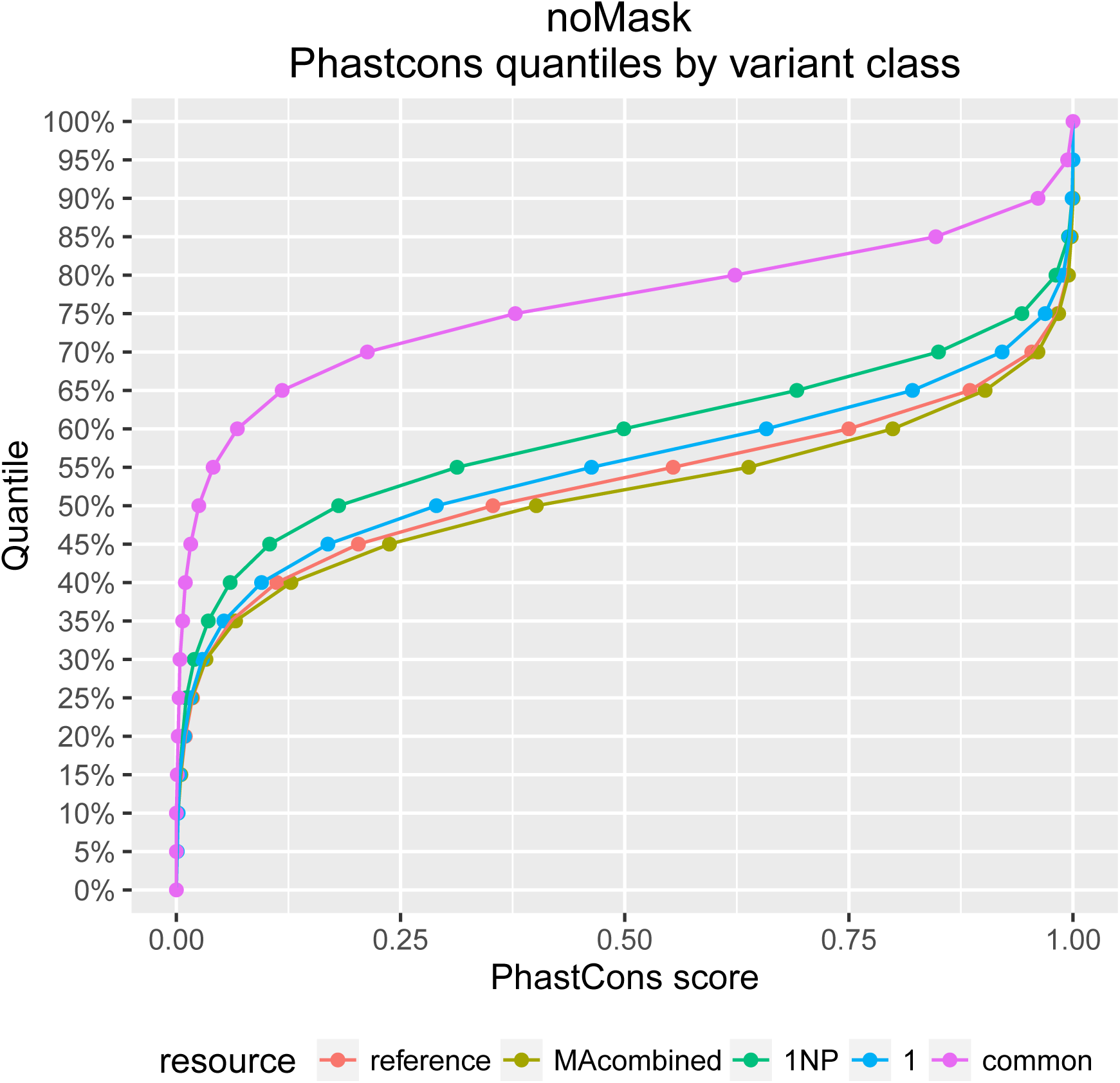
The empirical cumulative distirubution of phastCons scores for different classes of polymorphisms, and the reference genome.

**Figure SuppFigure-5:**
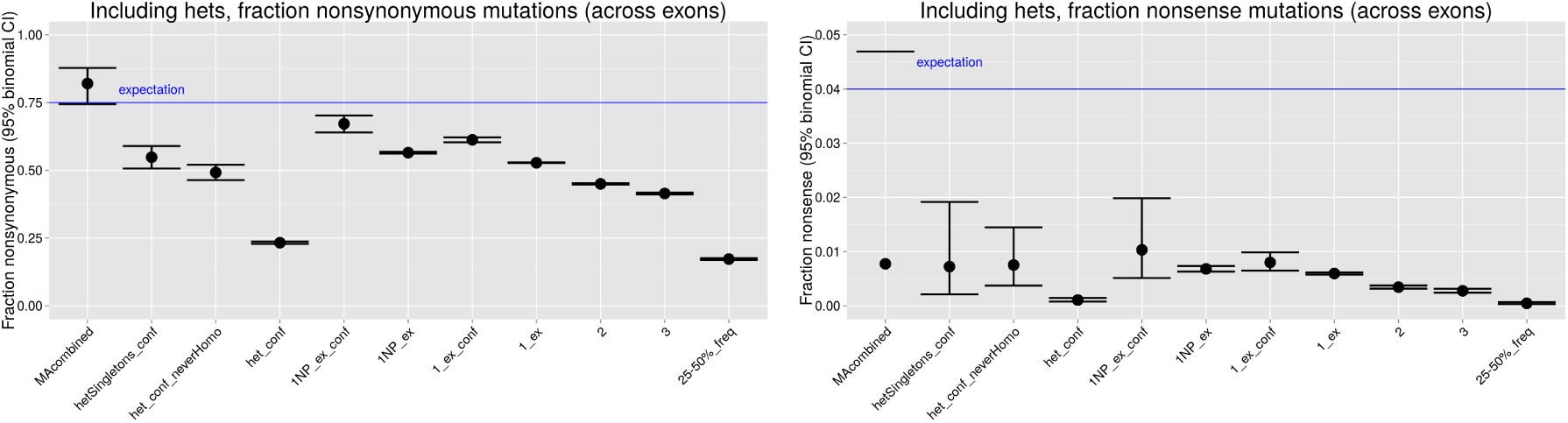
The fraction nonsynonymous (left) and nonsense (right) alleles present in heterozygous sites in the DGN. ‘hetSingletons_conf’ indicates a DGN singleton (frequency ~ 1/621) which was called as a heterozygote in DGN and confirmed heterozygous in the resequence data. ‘het_conf_neverHomo’ indicates all DGN polymorphisms (any frequency) which were called heterozygous in the DGN and also resequence data, and additionally were never found in a homozygous state in the DGN data. ‘het_conf’ indicates all DGN polymorphisms (any frequency) which were called heterozygous in DGN and also in the resequence data.

**Table SuppTable-7:** The results of a GO analysis of over-represented terms within conserved sites (phastCons=1), under-represented terms within sites containing common polymorphisms (frequency 0.2-0.5), and under-represented terms with sites containing rare polymorphisms (frequency ~ 1/621). See Methods for more detail on analysis.

|  | **Highly conserved sites**<br>**(i.e. phastCons = 1)** | **Common variants**<br>**(i.e. frequency 0.2-0.5)** | **Rare variants**<br>**(i.e. singletons)** |
| --- | --- | --- | --- |
| **GO test** | over-representation | under-representation | under-representation |
| **Interpretation of**<br>**GO test** | This reveals which functions tend to be highly conserved | This reveals which functions are depleted of common polymorphisms, suggesting new mutations tend to be weakly to strongly deleterious | This reveals which functions are depleted of rare variants, suggesting new mutations tend to be strongly deleterious |
| **GO categories**<br>**discovered** | **chromatin assembly or disassembly**<br>**nucleosome**<br>**nucleosome assembly**<br>**protein heterodimerization activity**<br>cytosol<br>DNA binding<br>translation<br>triplet codon-amino acid adaptor activity<br>ATP binding<br>cytoplasm<br>microtubule associated complex<br>neurogenesis<br>nucleus<br>plasma membrane<br>protein binding<br>regulation of transcription, DNA-templated<br>transcription factor activity, sequence-specific DNA binding | **chromatin assembly or disassembly**<br>**nucleosome**<br>**nucleosome assembly**<br>**protein heterodimerization activity**<br>cytosol<br>DNA binding<br>translation<br>triplet codon-amino acid adaptor activity<br>GGC codon-amino acid adaptor activity<br>heat shock-mediated polytene chromosome puffing | **chromatin assembly or disassembly**<br>**nucleosome**<br>**nucleosome assembly**<br>**protein heterodimerization activity** |

**Table SuppTable-8:** The results of G tests for triplet effect on the mutation spectrum, here for the singletons at frequency ~1/621. From left to right the columns refer to: variant dataset, triplet of interest (forward/reverse), mutation type 1, mutation type 2, mutation type 3, p value for G goodness of fit test (expected is total counts), corrected p value, and whether there is a significant effect of the triplet (p<0.01)

| ALTCOUNT | tripCombo | C->A/G->T | C->G/G->C | C->T/G->A | GtestPvalue | pCORR | pSIG |
| --- | --- | --- | --- | --- | --- | --- | --- |
| 1 | ACA/TGT | 962 | 557 | 1740 | 2.20E-02 | 3.50E-01 | FALSE |
| 1 | ACC/GGT | 538 | 293 | 952 | 3.50E-01 | 5.60E+00 | FALSE |
| 1 | ACG/CGT | 515 | 211 | 973 | 2.40E-03 | 3.90E-02 | FALSE |
| 1 | ACT/AGT | 670 | 421 | 1238 | 1.60E-03 | 2.60E-02 | FALSE |
| 1 | AGA/TCT | 754 | 455 | 1471 | 2.70E-02 | 4.40E-01 | FALSE |
| 1 | AGC/GCT | 924 | 469 | 1430 | 5.10E-05 | 8.20E-04 | TRUE |
| 1 | AGG/CCT | 680 | 321 | 1045 | 1.40E-03 | 2.20E-02 | FALSE |
| 1 | CCA/TGG | 1032 | 398 | 1538 | 2.00E-08 | 3.20E-07 | TRUE |
| 1 | CCC/GGG | 669 | 297 | 1111 | 5.00E-02 | 8.00E-01 | FALSE |
| 1 | CCG/CGG | 557 | 249 | 965 | 1.70E-01 | 2.80E+00 | FALSE |
| 1 | CGA/TCG | 492 | 302 | 1431 | 6.80E-20 | 1.10E-18 | TRUE |
| 1 | CGC/GCG | 698 | 329 | 1203 | 3.00E-01 | 4.80E+00 | FALSE |
| 1 | GCA/TGC | 1183 | 483 | 1676 | 3.60E-11 | 5.80E-10 | TRUE |
| 1 | GCC/GGC | 813 | 512 | 1733 | 2.10E-04 | 3.30E-03 | TRUE |
| 1 | GGA/TCC | 553 | 389 | 1573 | 7.80E-19 | 1.20E-17 | TRUE |
| 1 | TCA/TGA | 829 | 415 | 1728 | 1.20E-03 | 1.90E-02 | FALSE |
| 1 | AAA/TTT | 676 | 833 | 1138 | 6.00E-31 | 9.60E-30 | TRUE |
| 1 | AAC/GTT | 348 | 890 | 801 | 4.10E-02 | 6.60E-01 | FALSE |
| 1 | AAG/CTT | 354 | 579 | 986 | 1.60E-23 | 2.50E-22 | TRUE |
| 1 | AAT/ATT | 381 | 817 | 986 | 4.70E-04 | 7.60E-03 | TRUE |
| 1 | ATA/TAT | 216 | 836 | 787 | 8.80E-13 | 1.40E-11 | TRUE |
| 1 | ATC/GAT | 158 | 602 | 444 | 1.20E-10 | 1.90E-09 | TRUE |
| 1 | ATG/CAT | 225 | 714 | 583 | 3.60E-06 | 5.80E-05 | TRUE |
| 1 | CAA/TTG | 432 | 654 | 607 | 4.30E-14 | 6.90E-13 | TRUE |
| 1 | CAC/GTG | 376 | 694 | 540 | 1.60E-11 | 2.60E-10 | TRUE |
| 1 | CAG/CTG | 274 | 493 | 665 | 2.30E-06 | 3.70E-05 | TRUE |
| 1 | CTA/TAG | 146 | 397 | 482 | 1.20E-04 | 2.00E-03 | TRUE |
| 1 | CTC/GAG | 177 | 492 | 415 | 1.10E-02 | 1.70E-01 | FALSE |
| 1 | GAA/TTC | 248 | 635 | 482 | 1.70E-05 | 2.70E-04 | TRUE |
| 1 | GAC/GTC | 157 | 690 | 365 | 4.00E-28 | 6.50E-27 | TRUE |
| 1 | GTA/TAC | 148 | 605 | 504 | 9.40E-11 | 1.50E-09 | TRUE |
| 1 | TAA/TTA | 327 | 634 | 842 | 7.40E-07 | 1.20E-05 | TRUE |

**Table SuppTable-9:** The results of G tests for triplet effect on the substitution spectrum at four-fold degenerate sites. From left to right the columns refer to: evolved site dataset, triplet of interest (forward/reverse), mutation type 1, mutation type 2, mutation type 3, p value for G goodness of fit test (expected is total counts), corrected p value, and whether there is a significant effect of the triplet (p<0.01)

| tripID | tripCombo | C->A/G->T | C->G/G->C | C->T/G->A | GtestPvalue | pCORR | pSIG |
| --- | --- | --- | --- | --- | --- | --- | --- |
| fourD | ACA/TGT | 542 | 317 | 1614 | 9.40E-06 | 1.50E-04 | TRUE |
| fourD | ACC/GGT | 1557 | 989 | 2744 | 1.80E-37 | 3.00E-36 | TRUE |
| fourD | ACG/CGT | 853 | 515 | 1973 | 1.10E-01 | 1.80E+00 | FALSE |
| fourD | ACT/AGT | 626 | 447 | 2431 | 2.00E-26 | 3.20E-25 | TRUE |
| fourD | AGA/TCT | 428 | 276 | 1580 | 3.50E-16 | 5.70E-15 | TRUE |
| fourD | AGC/GCT | 436 | 284 | 2069 | 1.20E-49 | 1.90E-48 | TRUE |
| fourD | AGG/CCT | 866 | 555 | 2006 | 3.40E-02 | 5.50E-01 | FALSE |
| fourD | CCA/TGG | 1620 | 917 | 2693 | 1.00E-41 | 1.60E-40 | TRUE |
| fourD | CCC/GGG | 1338 | 844 | 2119 | 1.30E-50 | 2.10E-49 | TRUE |
| fourD | CCG/CGG | 1187 | 777 | 2754 | 3.60E-03 | 5.80E-02 | FALSE |
| fourD | CGA/TCG | 454 | 312 | 2016 | 8.40E-38 | 1.30E-36 | TRUE |
| fourD | CGC/GCG | 973 | 614 | 3095 | 2.00E-13 | 3.20E-12 | TRUE |
| fourD | GCA/TGC | 822 | 501 | 1930 | 2.30E-01 | 3.70E+00 | FALSE |
| fourD | GCC/GGC | 1195 | 767 | 2718 | 1.20E-03 | 1.90E-02 | FALSE |
| fourD | GGA/TCC | 906 | 558 | 2013 | 3.10E-03 | 5.00E-02 | FALSE |
| fourD | TCA/TGA | 588 | 408 | 2491 | 2.20E-39 | 3.40E-38 | TRUE |
| fourD | AAA/TTT | 143 | 364 | 201 | 9.20E-02 | 1.50E+00 | FALSE |
| fourD | AAC/GTT | 254 | 633 | 180 | 3.30E-28 | 5.20E-27 | TRUE |
| fourD | AAG/CTT | 233 | 464 | 450 | 5.10E-10 | 8.20E-09 | TRUE |
| fourD | AAT/ATT | 83 | 432 | 295 | 2.20E-09 | 3.60E-08 | TRUE |
| fourD | ATA/TAT | 67 | 323 | 144 | 2.40E-06 | 3.90E-05 | TRUE |
| fourD | ATC/GAT | 141 | 382 | 331 | 1.40E-04 | 2.20E-03 | TRUE |
| fourD | ATG/CAT | 93 | 460 | 185 | 6.80E-11 | 1.10E-09 | TRUE |
| fourD | CAA/TTG | 232 | 641 | 227 | 6.90E-16 | 1.10E-14 | TRUE |
| fourD | CAC/GTG | 354 | 1094 | 422 | 6.00E-19 | 9.50E-18 | TRUE |
| fourD | CAG/CTG | 407 | 942 | 689 | 3.00E-03 | 4.90E-02 | FALSE |
| fourD | CTA/TAG | 126 | 456 | 314 | 2.80E-03 | 4.50E-02 | FALSE |
| fourD | CTC/GAG | 451 | 729 | 948 | 4.80E-49 | 7.60E-48 | TRUE |
| fourD | GAA/TTC | 209 | 492 | 455 | 1.20E-07 | 1.90E-06 | TRUE |
| fourD | GAC/GTC | 423 | 1041 | 654 | 9.30E-02 | 1.50E+00 | FALSE |
| fourD | GTA/TAC | 111 | 502 | 188 | 3.60E-12 | 5.80E-11 | TRUE |
| fourD | TAA/TTA | 77 | 411 | 306 | 3.10E-11 | 5.00E-10 | TRUE |

**Figure SuppFigure-6:**
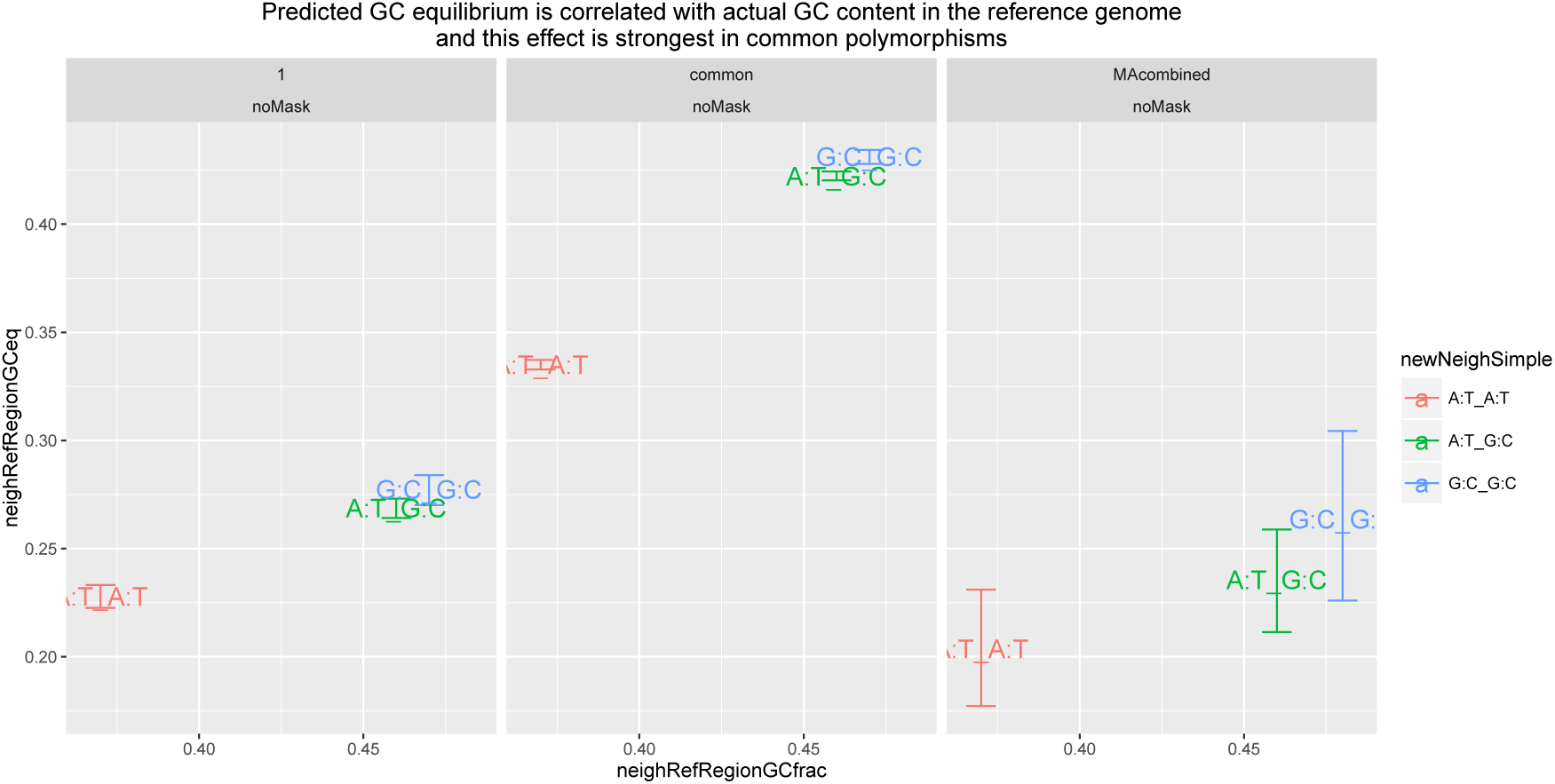
The predicted GC equilibrium (as calculated from MA mutations and DGN polymorphisms, y-axis) is correlated with the actual GC content of the reference genome (as calculated or the center base pair for each neighbor context, x-axis)

**Figure SuppFigure-7:**
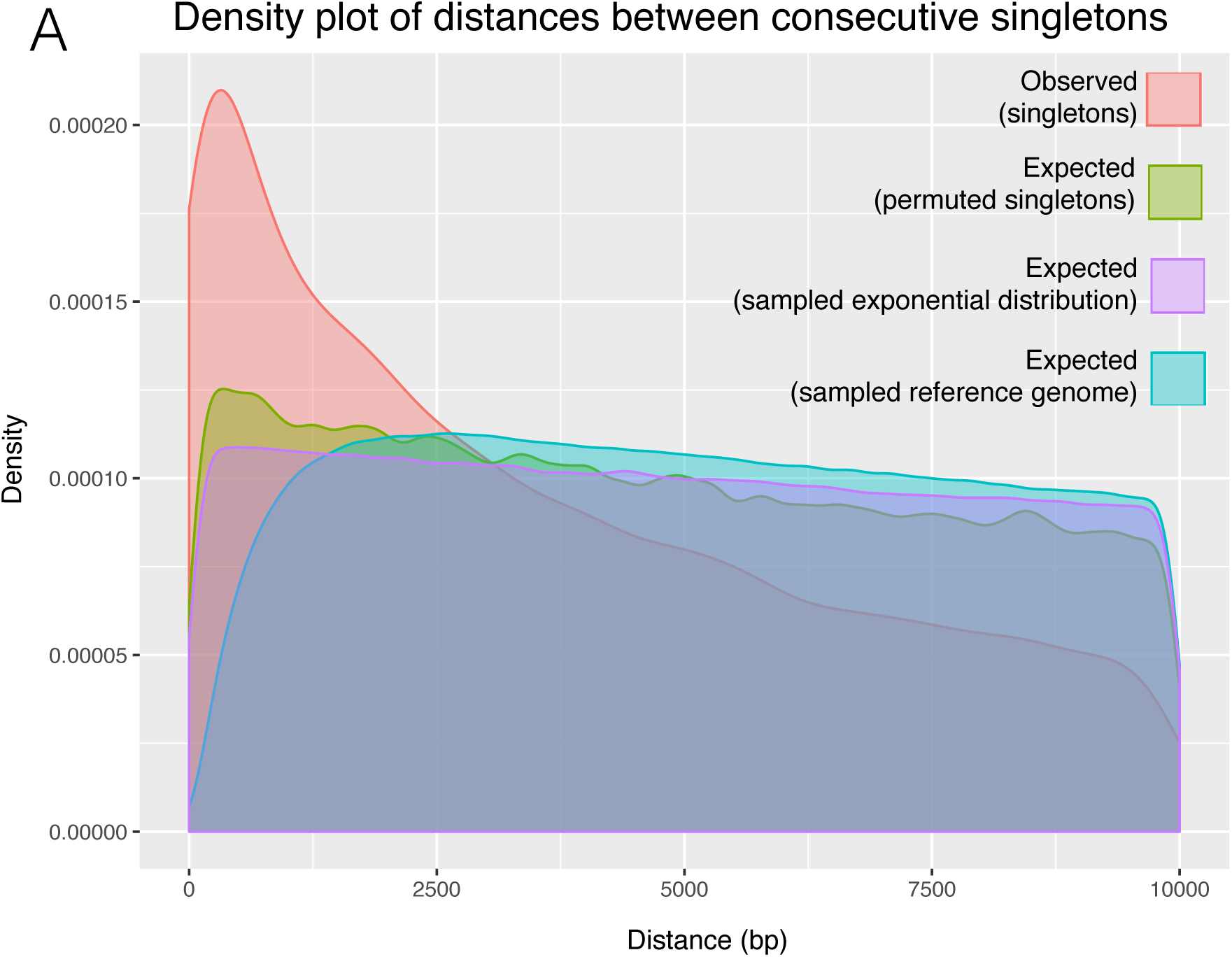
Density plots of sample data, and various possible expected distributions.

